# DNA methylation is a key mechanism for maintaining monoallelic expression on autosomes

**DOI:** 10.1101/2020.02.20.954834

**Authors:** Saumya Gupta, Denis L. Lafontaine, Sebastien Vigneau, Svetlana Vinogradova, Asia Mendelevich, Kyomi J. Igarashi, Andrew Bortvin, Clara F. Alves-Pereira, Kendell Clement, Luca Pinello, Andreas Gnirke, Henry Long, Alexander Gusev, Anwesha Nag, Alexander A. Gimelbrant

**Author notes:** Equal contribution (Saumya Gupta, Denis L. Lafontaine). Lead contact (AAG).

## Abstract

In diploid cells, maternal and paternal copies of genes usually have similar transcriptional activity. Mammalian allele-specific epigenetic mechanisms such as X-chromosome inactivation (XCI) and imprinting were historically viewed as rare exceptions to this rule. The discovery of mitotically stable monoallelic autosomal expression (MAE) a decade ago revealed an additional allele-specific mode regulating thousands of mammalian genes. However, despite its prevalence, the mechanistic basis of MAE remains unknown. To uncover the mechanism of MAE maintenance, we devised a small-molecule screen for reactivation of silenced alleles across multiple loci using targeted RNA sequencing. Contrary to previous reports, we identified DNA methylation as a key mechanism of MAE mitotic maintenance. In contrast with the binary choice of the active allele in XCI, stringent transcriptome-wide analysis revealed MAE as a regulatory mode with tunable control of allele-specific expression, dependent on the extent of DNA methylation. In a subset of MAE genes, allelic imbalance was insensitive to changes in DNA methylation, implicating additional mechanisms in MAE maintenance in these loci. Our findings identify a key mechanism of MAE maintenance, reveal tunability of this mode of gene regulation, and provide the essential platform for probing the biological role of MAE in development and disease.

## INTRODUCTION

In mammalian cells, the maternal and paternal gene copies tend to make an equal contribution to transcription (1). However, several allele-specific modes of gene regulation provide important exceptions. One such mode is genomic imprinting, where the allelic choice is determined by the parent of origin in about 200 mammalian genes (2). Another is X-chromosome inactivation (XCI), which randomly silences one of the two copies of the X chromosome in females (3), affecting over 800 X-linked genes. Additionally, olfactory sensory neurons express one allele of one out of ~1000 olfactory receptor genes (4).

The discovery of widespread monoallelic autosomal expression (MAE) greatly expanded our view of allele-specific gene regulation (5). Like XCI, MAE involves a random choice of the active allele during development, resulting in an epigenetic mosaic (6). Also like XCI, the allelic choice in MAE genes is mitotically stable; however, MAE genes can be expressed from both alleles in a subset of clonal lineages (**Fig.1A**). MAE had been observed in clonal populations of every cell type assessed (5, 7–13) and most MAE genes are highly cell-type specific (14). Cumulatively across cell types, an estimated 4000 human genes are subject to MAE (15), including genes implicated in cancer, neurodevelopmental disorders, and other diseases.

**Figure 1.**
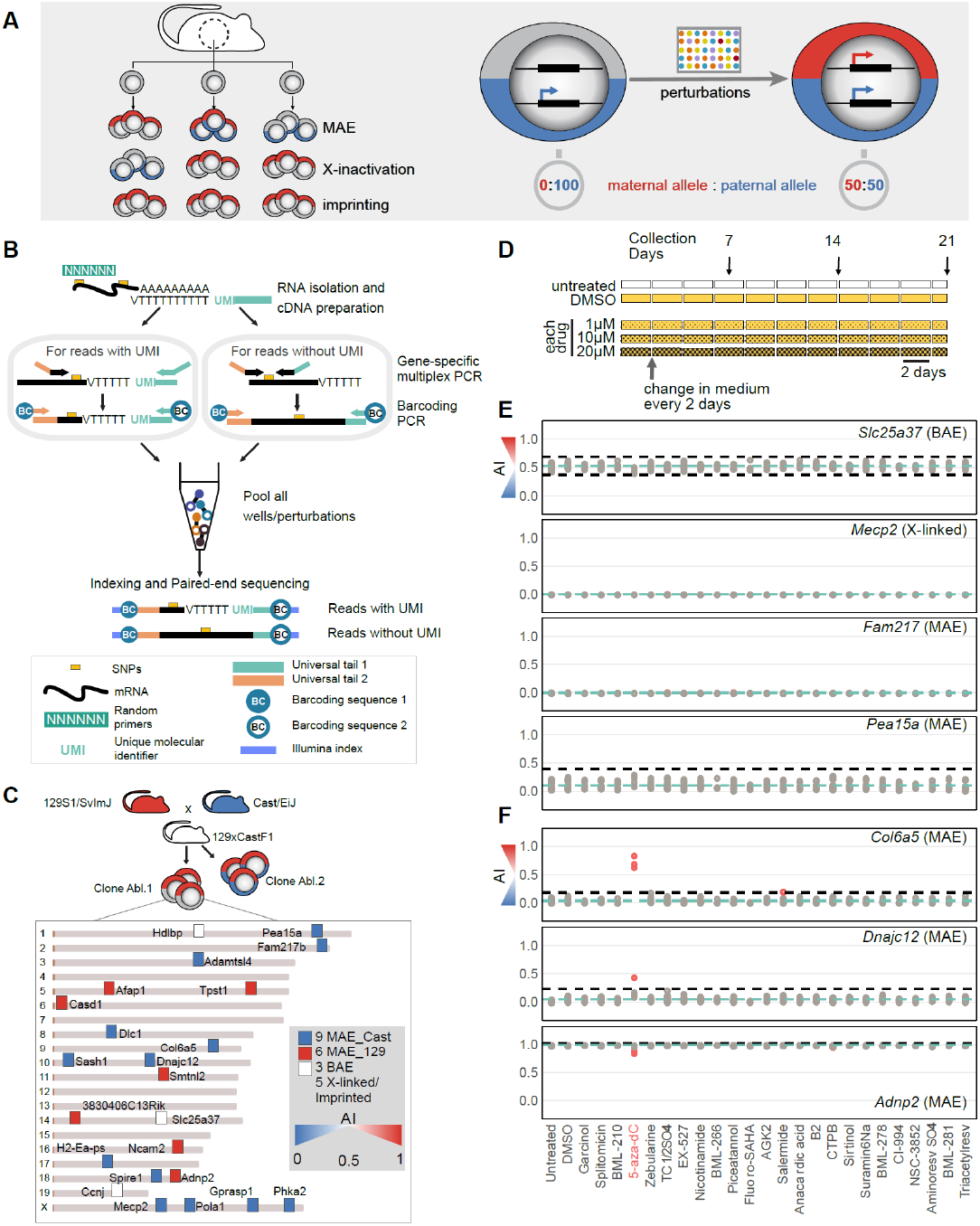
Perturbations that reactivate silenced alleles of genes with monoallelic expression identified using screening-by-sequencing. **(A)** Left: Different epigenetic modes of monoallelic expression. Note that while imprinting is uniform across cells, X chromosome inactivation and autosomal MAE result in epigenetic clonal mosaicism. Right: logic of the screen illustrated with a single locus in a clone with a completely silenced maternal allele. **(B)** Outline of Screen-seq methodology. Top to bottom: Cells are lysed in-plate, and in each well, RNA is isolated using SPRI beads. Two types of SNPs between parental genomes for the readout genes are targeted: those close to the poly-A tail enabling use of the UMI (*left*) and the rest that were targeted with two gene-specific primers with universal tails (*right*). Well-encoding is performed using primers targeting common adapters coupled with barcodes (BC1 and BC2). Then, all wells are pooled, Illumina sequencing adapters are added, and the pooled library is sequenced. See also **Suppl. Fig.S1** and **S2**. **(C)** 23 genes assayed in Screen-seq and their distribution in the mouse genome. Allelic imbalance (AI) of target genes in Abl.1 clone is reflected by the marker color. Centromeres (*brown*) on the left. **(D)** Drug treatment setup. Each of the tested 48 drugs were used in three concentrations: 1 μM, 10 μM and 20 μM in 1% DMSO. Fresh media and drugs were replaced every 2 days. Cells were collected on day 7, 14 and 21 and processed for Screen-seq. **(E, F)** Screen-seq results for a representative set of readout genes. All time and concentration points for a single drug shown in the same column. Each point shows allelic imbalance in one condition (AI=[129 counts]/[129 + Cast counts]). *Blue line*: mean AI for a gene across all conditions; *black dashed lines*: [Q1−3×IQR] and [Q1+3×IQR] (inter-quartile range); *red* points: outlier AI values (hits). **(E)** – genes showing no AI change in any condition. **(F)** – genes with significant changes in some conditions. Screen-seq results for all readout genes are in **Suppl. Fig.S3**.

The biological role of widespread MAE is presently unclear, though multiple lines of evidence imply it has a significant impact on organismal function. MAE has been shown to result in dramatic functional differences between otherwise similar cells; for example, the function of B cells in mice heterozygous for *Tlr4* depends on which allele is active in a given cell (16). Evolutionary and population-genetic analyses indicate conservation of the MAE status between human and mouse (13, 17), and selective advantage for individuals heterozygous for MAE genes (18, 19). The prevalence of cell surface molecules among proteins encoded by MAE genes prompted the hypothesis that MAE leads to increased variation in responses to extrinsic signals between otherwise similar cells (18).

Lack of knowledge about the underlying mechanism of MAE has severely limited research on MAE function. At present, no perturbation is known to affect the maintenance of allele-specific silencing in any MAE locus (20). MAE status correlates with histone modifications and DNA methylation in the gene body and putative regulatory sequences (8, 9, 12, 17, 21). However, inhibiting the DNA methyltransferases was reported to not affect the allelic imbalance in any of the tested MAE genes (8, 9), arguing against a mechanistic role of DNA methylation in MAE (20).

To understand the mechanistic basis of MAE, we devised a novel strategy for screening by targeted RNA sequencing and performed a small molecule screen for perturbations that affect allelic imbalance in expression in any of the 23 targeted genes across the mouse genome. We found that inhibition of methyltransferase *Dnmt1*-dependent DNA methylation reactivated silenced alleles in many MAE loci, showing that DNA methylation plays a major role in the MAE mitotic maintenance. At the same time, many MAE loci showed no significant changes in allele-specific expression upon DNA demethylation, suggesting that MAE is mechanistically heterogeneous. Application of a highly stringent statistical approach to transcriptome-wide differential analysis of allele-specific expression (22) revealed the existence of multiple mitotically stable states of allelic imbalance, which correlated with the extent of DNA methylation. We conclude that DNA methylation acts as a fine-tuning mechanism for MAE loci, controlling allele-specific transcription as a rheostat, as opposed to an on-off switch.

## RESULTS

### Screening-by-sequencing approach for sensitive detection of allele-specific expression

To screen for reactivation of a silenced allele, we looked for shifts in allelic imbalance (AI, the fraction of one allele over the total allelic counts; **Fig.1A**, right) upon drug treatment. In order to increase the likelihood of detecting AI shifts among genes with potentially different regulation, our screening approach would ideally combine the ability to assess multiple readout genes, sensitivity to AI changes, and the throughput to process multiple samples after exposure to an array of perturbations.

We designed a screening-by-sequencing strategy, Screen-seq, to satisfy all of these requirements. In cells with heterozygous genomes, allele-specific expression can be assessed without the need for any engineered reporters and by relying on the detection of single nucleotide polymorphisms (SNPs). Precision and sensitivity of the AI measurement in RNA sequencing critically depend on the depth of SNP coverage (22). Sequencing of SNP-containing amplicons from multiplexed RT-PCR as the readout allows for very deep coverage and thus a highly precise AI measurement.

The experimental flow of Screen-seq is outlined in **Fig.1B**. Cells were grown and lysed in 96-well plates; RNA isolated using magnetic beads, and cDNA synthesized with a mix of random primers and oligo-dT primers with Unique Molecular Identifiers [UMIs, (23, 24)]. This mix allowed targeting of two types of SNPs in the next step, multiplex PCR: SNPs close to the 3’-end enable the use of oligo-dT-UMIs followed with a gene-specific primer, while other SNPs were targeted with two gene-specific primers in random-primed cDNA. Next, plate- and well-encoding barcodes were added using PCR. The reactions from all the wells were pooled, Illumina adaptors added, and the pooled library was sequenced. Finally, SNP counts were assigned to specific genes, and barcodes to specific plates and wells with a specific perturbation.

To allow analysis of MAE genes, which show different AI in different clones, we performed our screen in a monoclonal line of pro-B cells (Abl.1). We have previously characterized allele-specific expression in several such clones, including Abl.1 and other clones used in this study (13, 14). These cells were derived from a female 129S1/SvImJ × Cast/EiJ F1 mouse, then immortalized using the Abelson murine leukemia virus (25) and cloned through single-cell sorting. In this mouse cross, the median distance between SNPs in the non-repetitive genome is ~80 bp and almost all cDNAs have at least one informative SNP.

For readout, we selected 27 SNPs in 23 target genes across the genome, including 15 clone-specific MAE genes as well as three biallelic, one imprinted and four X-inactivated loci (**Fig.1C, Suppl. Table S1**). The selected MAE genes showed AI>0.9 or AI<0.1 in the Abl.1 clone (AI=[129 counts]/[129 + Cast counts]), while showing opposite bias or biallelic expression in another clone, Abl.2 (13, 14). Targeted MAE genes spanned a range of expression levels and extent of allelic bias in the screening clone, Abl.1; some showed complete silencing of one allele (such as *Afap1*, AI = 1.0), while others showed strong but incomplete bias (such as *Dlc1*, AI = 0.1).

We first tested that these assays were able to detect changes in AI. Since no perturbations are known that can change AI in any locus, much less in all targeted loci, for the control experiments we titrated known mixes of genomic DNA from liver tissue of the parental mouse strains, 129S1/SvImJ and Cast/EiJ. Expected and measured AI were highly concordant (R^2^ ≥ 0.99) at >1000 reads/SNP (**Suppl. Fig.S1**). We also compared AI sensitivity for UMI and non-UMI assays, by designing both types of assays for a subset of genes where the position of SNPs allowed that. For this, we used mixes prepared from total RNA from the spleens of the mice of the parental mouse strains. AI measurements were highly concordant between the UMI and non-UMI assays (R^2^ ≥ 0.97, **Suppl. Fig.S2)**.

Based on these pilot experiments, we concluded that Screen-seq can be used for sensitive detection of AI changes in the targeted loci.

### Identification of perturbations that affect allele-specific gene expression

Clone-specific MAE has been associated with specific chromatin signatures, i.e., combinations of histone modifications in human and mouse cells (8, 12, 14), suggesting that chromatin modifying mechanisms might be involved in MAE maintenance. We thus assessed the impact on AI in the targeted loci of treatment with a set of 43 small molecules with known effects on the activity of the enzymes involved in the deposition and removal of methylation and acetylation marks on histones and DNA (**Suppl. Table S3**). Abl.1 cells in 96-well plates were exposed for 21 days to individual drugs in regular growth conditions (**Fig.1D**). Each drug was applied in three final concentrations (1 μM, 10 μM and 20 μM in 1% DMSO). Controls were untreated cells and cells with only solvent (1% DMSO) added. Fresh media (with or without drugs, as appropriate) was replaced every two days. On days 7, 14, and 21, aliquots of cells were removed for analysis.

For 19 of the 43 drugs, no live cells were evident after six days, at any drug concentration. Each cell collection thus involved only 72 wells with treated cells (24 remaining drugs at three concentrations) and 24 wells with controls (12 untreated and 12 vehicle-treated cells). Taken together, in this Screen-seq experiment we assessed 7,776 experimental points (allele-specific measurements of 27 SNPs in 23 genes × 96 wells × 3 time points).

With a targeted RNA-seq library, only a very moderate amount of sequencing was needed to reach the coverage depth required for sensitive allele-specific analysis. At 1,000 reads per experimental point, fewer than 10×10^6^ sequenced fragments were needed for the entire screen.

As potential hits, we identified conditions resulting in outlier AI values (**Fig.1E,F**; see Methods for details). AI measurements were highly uniform for some genes (e.g. *Fam27b* or *Mecp2*) across drug concentrations and time points, while there was more variation in other genes (e.g., *Pea15a* or *Col6a5*). To allow for variation in assay sensitivity, each readout gene was analyzed independently of the rest. Outliers were identified using highly stringent criteria (see Methods).

As expected for stably maintained allele-specific expression, in the untreated cells there were no outliers for any of the readout genes. The most pronounced outliers (red in **Fig.1F**) were observed for 3 MAE readout genes in the presence of 5-aza-2’-deoxycytidine (5-aza-dC). There were also significant AI shifts in single reporter genes after exposure to histone deacetylase modulators Salermide and BML-278 (complete Screen-seq results are in **Suppl. Fig.S3** and **Suppl. Table S4**). The magnitude of the observed shifts varied between genes and conditions, including drug concentration and exposure times. The most striking example is a shift in *Col6a5* gene from baseline AI≈0.1 baseline in the control to AI≈0.8 after 7 days in the presence of 1 μM 5-aza-dC (**Fig.1F**). More subtle, significant shifts were observed for the MAE genes *Adnp2* (from AI=1.0 to AI=0.8) and *Dnajc12* (AI≈0.1 to AI≈0.2). Notably, in other tested genes, no AI shift was observed in 5-aza-dC (**Suppl. Fig.S3)**. We focused on characterizing the strongest primary hit, 5-aza-dC.

### 5-aza-dC affects allele-specific expression of autosomal MAE genes via DNA demethylation

To validate the candidate hit 5-aza-dC, a classic DNA demethylation agent (26), we performed several sets experiments. First, we took advantage of the fact that the Screen-seq protocol leaves enough RNA and cDNA for re-testing. We measured AI in the same samples using an orthogonal method, droplet digital PCR (ddPCR, a highly sensitive approach to measuring allelic frequencies (27)). In addition to using a different readout method, we assessed different SNPs than those used for Screen-seq for the same genes. Using cDNA from cells treated with 1, 10 and 20 μM of 5-aza-dC for 7 days, we performed ddPCR to assess reactivation of the silenced maternal allele of the *Col6a5* and *Dnajc12* genes. Confirming the results from Screen-seq, ddPCR measurements showed a similarly striking shift in *Col6a5* AI from a paternal bias to maternal bias (AI=0.1 to 0.8) after 7 days in 1 μM 5-aza-dC (**Fig.2A,B**). Also confirming the Screen-seq results, AI for *Dnajc12* gene showed relaxation towards a more biallelic expression, with AI shifting from 0 to 0.1 in 1 μM 5-aza-dC and to 0.3 in 20 μM 5-aza-dC in 7 days (**Fig.2B**).

**Figure 2.**
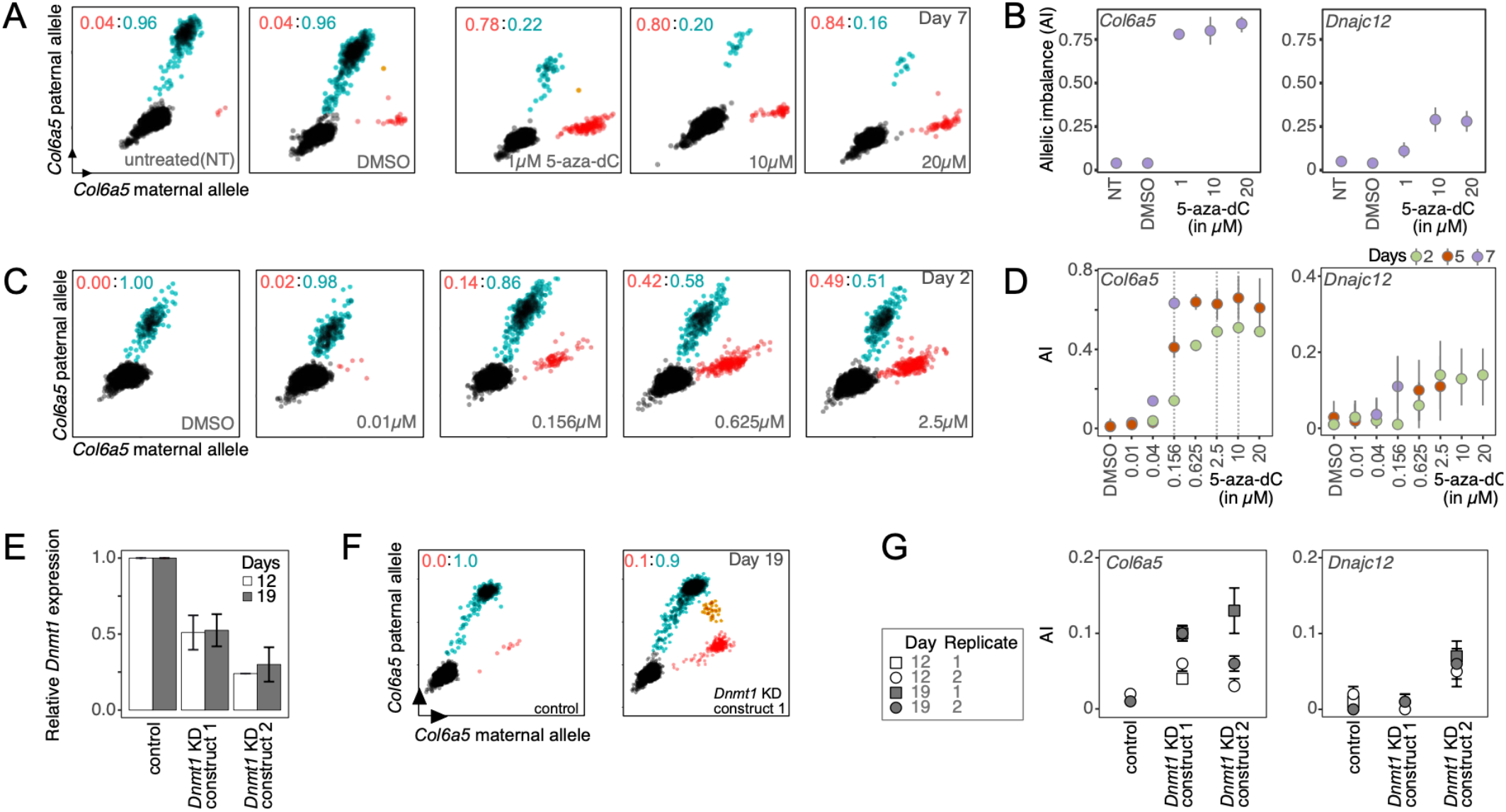
Hit validation for 5-aza-2’-deoxycytidine. **(A, B)** Confirmation of Screen-seq results for 5-aza-2’-deoxycytidine (5-aza-dC) treated cells, using an orthogonal method to measure AI. cDNA samples from day 7 of screening were assessed using droplet digital PCR (ddPCR) with allele-specific fluorescent probes. **(A)** Scatterplots for 20,000 droplets targeting the readout gene, *Col6a5*. 5-aza-dC concentration is shown in the plots. *Black*: empty droplets; *blue*: droplets with the Cast paternal allele amplified (labeled by FAM fluorophore); *red*: droplets with the 129 maternal allele amplified (labeled by HEX fluorophore). Ratio of red:blue droplets shown. Red value is AI as used throughout the manuscript. **(B)** *Left:* Summary of AI measurements shown in (A) for *Col6a5*; *right:* summary of AI measurements for *Dnajc12*. **(C, D)** Biological replicate of Abl.1 cells were treated with 5-aza-dC and AI was measured using ddPCR. **(C)** – Scatterplots as shown in (A) after 2 days of exposure. **(D)** – summary of AI measurement for *Col6a5* (left) and *Dnajc12* (right) after 2, 5, and 7 days of exposure. Grey vertical dashed lines for *Col6a5* dose-response were used to determine “low”, “medium” and “high” 5-aza-dC concentrations for the genome-wide experiments. Results for readout gene *Adnp2* are in **Suppl. Fig.S4**. See also **Suppl. Fig.S5-7. (E, F, G)** Analysis of *Dnmt1* knock-down (KD) in Abl.1 cells. **(E)** Real-time quantitative PCR (RT-qPCR) analysis of *Dnmt1* relative expression (expression in the empty vector control, normalized to *Nono*, taken as 1.0). Abl.1 cells were transduced with an empty plKO vector (control) or with 2 separate *Dnmt1* shRNA knockdown constructs (*Dnmt1* KD construct 1 or 2) and grown for 2 days. Transduced cells were then selected by growing in the presence of a selection antibiotic for an additional 17 days. RT-qPCR quantification was performed on cells collected 19 days after transduction. Mean and S.E.M for 3 technical replicates are shown. **(F)** Representative scatterplots show AI measurement for *Col6a5* in the transduced Abl.1 cells. AI was measured using ddPCR. **(G)** Summary of the AI measurement for *Col6a5* (left) and *Dnajc12* (right) after *Dnmt1* KD.

In biological replicate experiments, the Abl.1 clonal cells were exposed to a range of concentrations of 5-aza-dC for varying times. Using ddPCR, we observed that the maternal allele of *Col6a5* was reactivated in a dose- and time-dependent manner (**Fig.2C,D**). AI shifts for *Dnajc12* and *Adnp2* were also concordant with those observed in Screen-seq (**Fig.2D** and **Suppl. Fig.S4**). Taken together, these observations show that 5-aza-dC causes a shift in allelic imbalance in a subset of MAE genes.

A closely related compound, 5-aza-cytidine (5-aza-C), is also a well-known demethylating agent, although less potent and toxic than 5-aza-dC (28). Since 5-aza-C was not one of the perturbagens tested in our screening, we assessed whether it had a similar effect as 5-aza-dC on AI changes. Within 2 days of treatment with 10 μM 5-aza-C, the AI of *Col6a5* shifted from 0 to 0.2, and to 0.6 after 5 days in 2 μM 5-aza-C (**Suppl. Fig.S5**). Another MAE readout gene, *Dnajc12*, showed a shift in AI from 0 to 0.1 within 2 days in 2 μM 5-aza-C. This further supports the role of DNA methylation in MAE maintenance.

5-aza compounds at high concentrations are cytotoxic and cause cell cycle arrest (29). We asked whether shifts in AI in the target genes might be due to nonspecific cytotoxicity. In the presence of 2% DMSO, higher than the 1% concentration used as a drug solvent, the Abl.1 clonal cells viability was reduced to 34% after 2 days, similar to their viability after 5 days in 2.5 μM 5-aza-dC (**Suppl. Fig.S6**). In contrast to the AI shifts in the presence of 5-aza-dC and 5-aza-C (**Fig.2D** and **Suppl. Fig.S5**), no changes in AI were observed for the MAE readout genes, *Col6a5* and *Dnajc12*, in 2% DMSO (**Suppl. Fig.S7**), indicating that AI shifts are not a generalized feature of cells under stress.

To test if the effect of 5-aza-dC on allele-specific expression was specific to inhibition of methyltransferase activity, we assessed changes in AI in response to the knock-down of *Dnmt1*, the main maintenance methyltransferase in mammals (30). Abl.1 cells transduced with *Dnmt1* shRNA constructs showed 2-fold and 4-fold decrease in *Dnmt1* RNA abundance (**Fig.2E**), and the corresponding partial reactivation of silenced alleles of *Col6a5* and *Dnajc12* (**Fig.2F,G**).

Taken together, these observations indicate that *Dnmt1*-dependent DNA methylation is a molecular mechanism involved in AI maintenance for at least some MAE genes.

### Changes in allelic imbalance are long-term and rheostatic

The shifts in AI we observe could be consistent with two different mechanisms. The shift could result from the long-term change in the mitotically stable state of allele-specific gene regulation. Alternatively, it could be due to short-term changes, e.g., because of stress caused by drug exposure. We thus asked if the changes in AI were mitotically stable and enabling long-term maintenance, the hallmark of autosomal MAE.

To address this question, we performed a treatment-and-recovery experiment (**Fig.3** and **Suppl. Fig.S8A**). First, Abl.1 cells (with doubling time of ~12 hrs) were exposed to 5-aza-dC; after two days, cells were washed and incubated further in the regular growth medium. **Fig.3A and 3B** show the AI readout for *Col6a5* gene (similar results were seen with *Dnajc12* gene, **Suppl. Fig.S8B**). After two days of treatment and three days of recovery, AI reached levels that remained stable through days 9 and 12. This shows that AI shifts resulting from 5-aza-dC treatment were maintained over multiple subsequent cell divisions. Such stability is consistent with DNA methylation as the molecular mechanism that maintains the long-term memory of AI state of MAE genes in clonal cells. A continuing AI shift over the first three days of recovery is consistent with the cell population right after treatment being heterogeneous and containing some remaining fraction of cells with the readout gene in the initial state of AI≈0. By day 5, that fraction would be replaced by cells in the new stable state of DNA methylation, and the new state would then be maintained through days 9 and 12.

**Figure 3.**
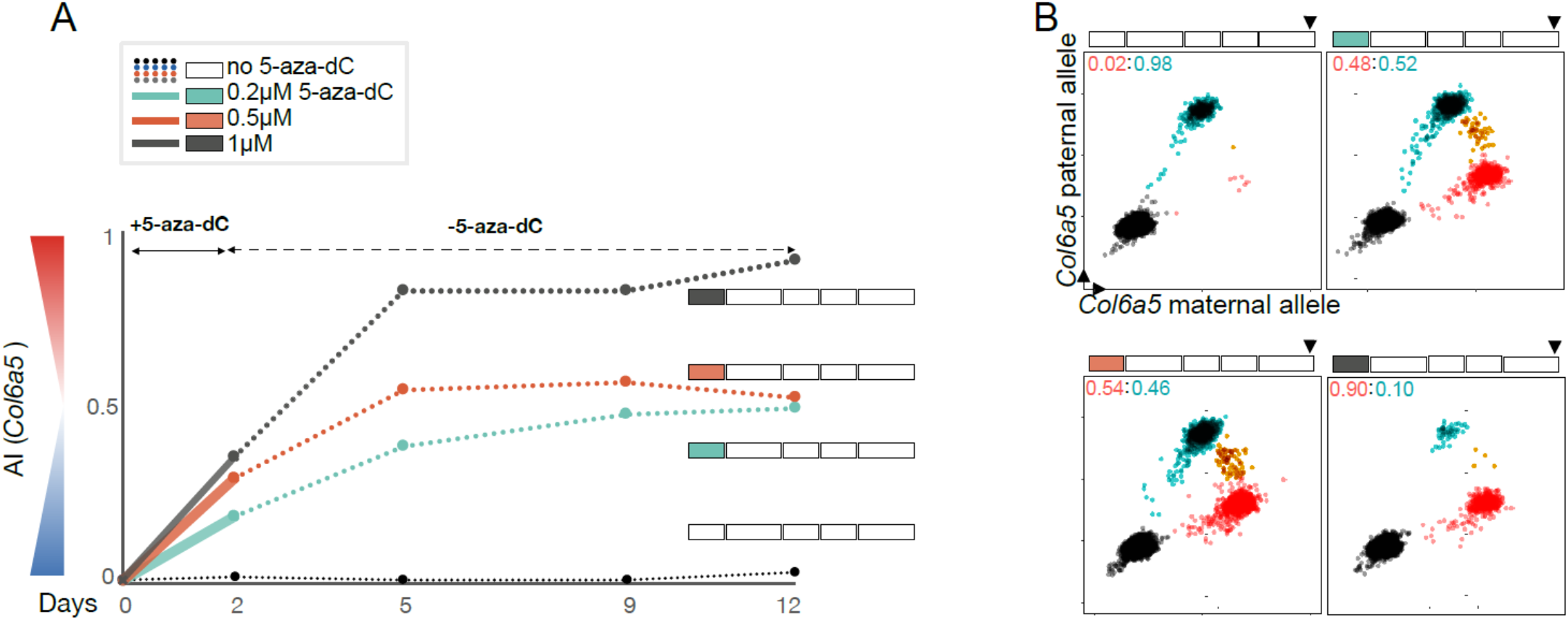
Long-term changes in the mitotic memory of allelic imbalance after exposure to 5-aza-dC and recovery. **(A)** 5-aza-dC exposure/recovery experiment in Abl.1 cells. Drug treatment setup is shown next to the line-plots (*small box*-2 days, *long box* - 3days). Media changes are shown as breaks in the boxes in the drug treatment setup. Cells were exposed to 0.2, 0.5 or 1 μM 5-aza-dC (denoted by colors) in growth medium. Cells were moved to the medium without any drug after 2 or 5 days, and collected for analysis at days 2, 5, 9, and 12. AI measurements for *Col6a5* across time points using ddPCR are summarized in the line-plot. Results shows here are for cells that were exposed to 5-aza-dC for 2 days. See **Suppl. Fig.S8** for the complete results. **(B)** ddPCR scatterplots for *Col6a5* on day 12 (shown by arrows on the drug treatment setup) after recovery [as summarized in **(A)**].

We observed in other experiments (see **Fig.2**) that the extent of allelic shift was dose dependent. Notably, after recovery, the eventual stable AI states were also dependent on the 5-aza-dC concentration during cell exposure (**Fig.3**). This shows that 5-aza-dC-dependent allele-specific regulation acts not as an on-off switch, but rather as a rheostat, with multiple stable intermediate states.

Similarly, the extent of the AI shift correlated with the length of drug exposure. After two days of exposure, half the cells were moved into the regular growth medium, while the other half were exposed to the same drug concentration for an additional 3 days. Additional exposure led to further AI shifts (**Suppl. Fig.S8A**). Note that longer exposure to 5-aza-dC decreased cell viability to the point where insufficient number of viable cells were present after day 5 for reliable analysis (**Suppl. Fig.S8C**). The same eventual AI shifts were reached by cells after 5 days of treatment as after treatment and recovery: AI≈0.9 was reached after (i) two days of treatment with 1 μM 5-aza-dC with three days recovery; (ii) five days treatment with 0.5 μM; and (iii) five days treatment with 1 μM (**Suppl. Fig.S8B**). This is consistent with a regulatory locus reaching complete demethylation in all conditions.

### Genome-wide allele-specific impact of DNA demethylation

We assessed the global impact of 5-aza-dC on allele-specific DNA methylome and transcription by exposing Abl.1 cells for 2 days to low (0.2 μM), medium (2 μM) or high (10 μM) concentrations of 5-aza-dC, compared with solvent only (1% DMSO) as the control, and performing RNA and reduced-representation bisulfite (RRBS (31)) sequencing. When not resolving allele-specific signal, both RNA abundance and the extent of DNA methylation behaved as expected. DNA methylation levels substantially decreased in the presence of 5-aza-dC (**Suppl. Fig.S9**). Consistent with the overall transcriptional derepression due to DNA demethylation, RNA abundance generally increased (**Suppl. Fig.S10**).

To resolve allele-specific signal in transcription, we applied Qllelic, a novel, highly stringent approach to the analysis of allele-specific expression, which uses replicate RNA-seq libraries to account for technical AI overdispersion and to minimize false positives for differential AI (22). Note that accounting for the whole-transcriptome scale of RNA-seq necessarily made some AI changes, readily detectable in targeted assays, fall short of statistical significance in RNA-seq. For example, of the three genes with shifts in 5-aza-dC detected with Screen-seq (**Fig.1**) and confirmed using ddPCR (**Fig.2**), only the shift in *Col6a5* reached statistical significance in the RNA-seq analysis (**Suppl. Table S6**). This underscores the utility of targeted assays for screening purposes.

Applying this stringent analysis to the Abl.1 RNA-seq data, we saw 51 genes with a significant AI shift after exposure to the low 5-aza-dC concentration, 145 genes in medium, and 140 genes in high 5-aza-dC (**Fig.4A**, **Suppl. Fig.11** and **Suppl. Table S7**). The direction of AI shifts at different concentrations of 5-aza-dC all agreed, with higher concentration generally corresponding to larger shift (**Fig.4C**). No known imprinted genes showed AI changes under these conditions, and for the 3 X-linked genes with statistically significant changes, the absolute shift was very small (e.g., shift in *Hccs* from AI=0.0 to 0.05, **Suppl. Table S7**), suggesting more robust mitotic maintenance of imprinting and X-inactivation than MAE.

**Figure 4.**
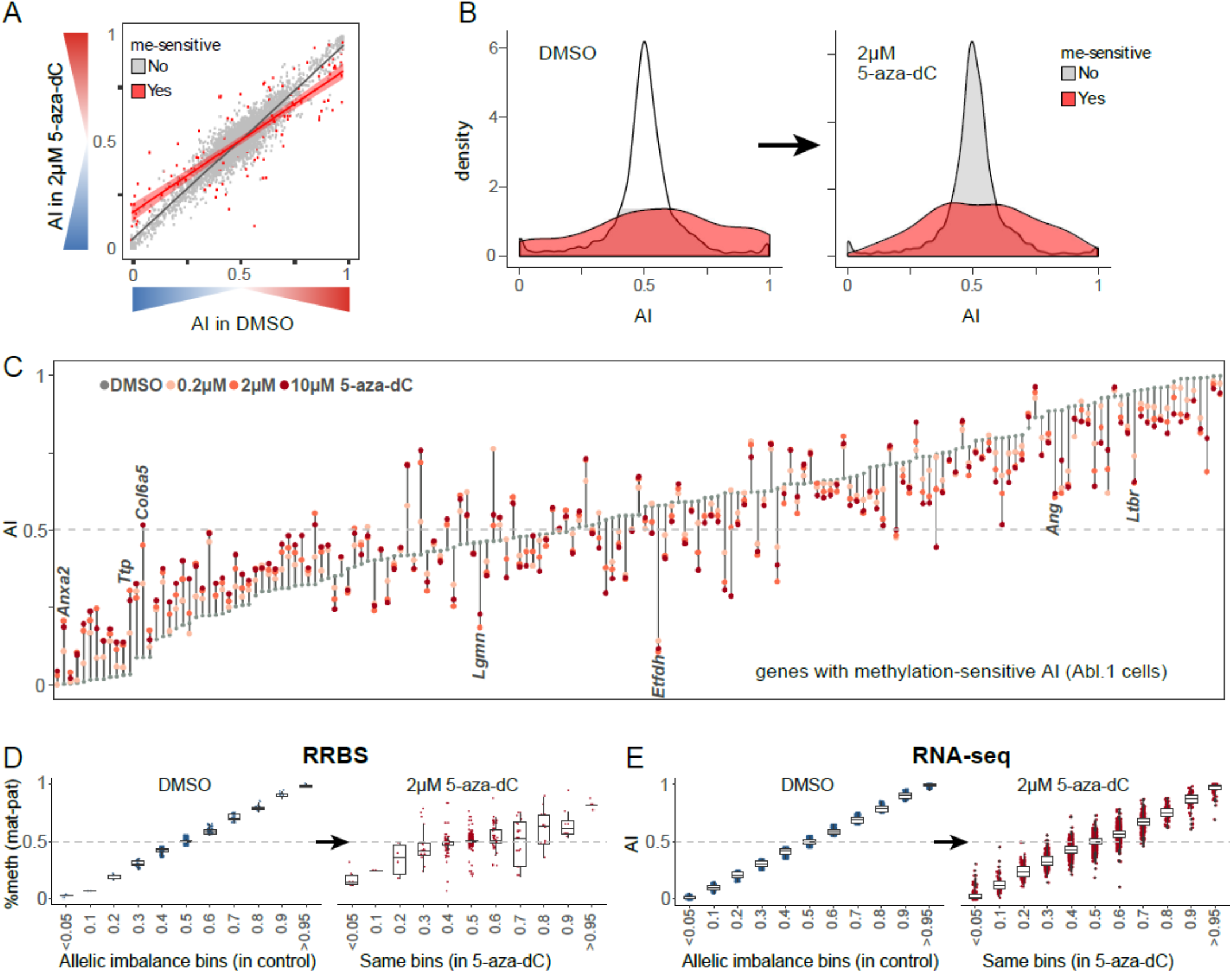
Genome-wide allele-specific effects of cell exposure to 5-aza-dC. **(A)** Comparison of allele-specific expression in Abl.1 cells in control cells and 2 μM 5-aza-dC. Genes with significant shift in AI [methylation-sensitive genes] are shown in *red*. For other 5-aza-dC concentrations, see **Suppl. Fig.S11**. **(B)** Density plots for distribution of AI values of genes with no significant changes in AI (*grey*) and with changes (*red*) [same experiment as in **(A)**]. Note that grey and red areas are plotted to be equal. Left: control; right: 2 μM 5-aza-dC. For other concentrations, see **Suppl. Fig.S19**. **(C)** Shifts of AI in expression for methylation-sensitive genes [same experiment as in **(A)**]. Genes shown in the order of their nominal AI in control (*grey circles*). AI after treatment is shown (color denotes 5-aza-dC concentration). For other clonal lines, see **Suppl. Fig.S15.** For full list of methylation-sensitive genes, see **Suppl. Table S7** and **Suppl. Fig.S12.** **(D-E)** Allele-specific changes in DNA methylome and transcriptome in Abl.1 cells. **(D)** Allele-specific DNA methylation analysis of RRBS (reduced representation bisulfite sequencing) data. *Left:* Sites were binned by AI, defined as difference between methylated fraction of CpGs on maternal and paternal alleles in control conditions (1% DMSO). *Right:* AI in the same bins after treatment. See also **Suppl. Fig.S9**. **(E)** Genes were binned by AI values, in control (*left*) and after treatment (*right*). Note: in (D,E), AI bins are labeled with central value X, with the bin covering the range of (X−0.05; X+0.05]. See also **Suppl. Fig.S10** and **S13** for bins showing significant shifts between control and treatment.

Known examples of changes in transcription AI, such as loss of imprinting and loss of X-inactivation in cancer (32, 33) involve relaxation of very strong allelic biases towards AI=0.5. This aligns with the notion of genetic variation by itself making a small contribution to allelic bias, while epigenetic mechanisms can impose dramatic allelic imbalance. In Abl.1 cells, observed initial AI values and shifts were following this pattern in some genes (e.g., *Anxa1, Ttp, Ang*, and *Ltbr;* **Fig.4C**)).

Use of replicate RNA-seq libraries and Qllelic enabled highly confident estimation of AI, revealing that most of the genes with AI shifts had initial (before treatment) AI values between extreme bias and AI=0.5, with these intermediate values much more common among these genes than in the transcriptome as a whole (**Fig.4B**). In addition, the direction of the AI shifts was also often unexpected. Out of 145 genes with significant shifts in AI, 44 genes (30%), including *Lgmn* and *Etfdh,* showed greater allelic bias (further away from AI=0.5) after 5-aza-dC exposure than in their initial state (at 2 μM; 13/51 genes (26%) at 0.2 μM, and 37/140 (26%) at 10 μM; **Fig.4C**).

To analyze changes in allele-specific signal in the DNA methylome from RRBS data, we first grouped sites by AI bins, since very few individual sites had sufficient coverage for statistical significance in a genome-wide analysis (**Suppl. Table S9**). In this analysis (**Fig.4D**), every bin with significant AI changes showed a shift towards AI=0.5 (**Suppl. Fig.S13A**). This appears inconsistent with ~30% of the genes showing increased AI in expression after exposure to 5-aza-dC (**Suppl. Fig.S13B** and **Fig.4C**). Furthermore, using similar binning of genes by AI, although the overall relaxation towards AI=0.5 was statistically detectable, the extent of this AI shift was much less pronounced than for RRBS (**Fig.4E**).

Taken together, these observations show that AI in transcription is not necessarily a simple predictor of AI in DNA methylation. Instead, allele-specific gene expression is often determined by an interplay of genetic and epigenetic regulation.

### DNA demethylation leads to increased similarity between clones

An MAE gene can show extreme allelic bias in one clone and biallelic expression in another; this is a distinct feature of MAE genes, in contrast to XCI and imprinting (5, 34), and it directly contributes to clonal heterogeneity. In previous RNA-seq studies of MAE, assignments of allelic states were categorical and based on arbitrary thresholds. Gene expression would be classified as “monoallelic” if the major allele constitutes over 85% (9), or 66% (12), or 98% (7).

We took advantage of the precision of AI estimates from Qllelic to perform differential analysis of allele-specific expression across several clones and its changes after DNA demethylation. In addition to Abl.1 clone, we assessed allele-specific expression in clones Abl.2, Abl.3, and Abl.4, all derived from 129xCastF1 mice (13) and thus genetically nearly identical. To control for possible loss of heterozygosity events, we performed exome sequencing and removed from comparisons any locus with pronounced allelic bias in genomic DNA (AI<0.3 or AI>0.7; see Methods).

To compare shifts in AI after DNA demethylation, we first assessed the toxicity of 5-aza-dC across clones. The viability of Abl.2, Abl.3, and Abl.4 clones was affected at 0.2 μM similarly to the Abl.1 clone at 2 μM (**Suppl. Fig.S14**). After two days’ exposure to these equitoxic concentrations, 677 genes between the four assessed clones showed AI shifts: 252 genes in Abl.2, 282 in Abl.3, and 172 in Abl.4 cells (**Suppl. Table S7**). AI shifts had similar features in all clones. Initial and final AI states of the affected genes were distributed similarly to Abl.1 clone, and the direction of shifts was also similar (**Suppl. Fig.S15**).

There were 1,767 genes with significant differential AI between clones (**Suppl. Table S11**), and 346 (20%) of these showed significant changes of AI after DNA demethylation (**Suppl. Table S12**). AI in the other genes with differential AI did not change after exposure to 5-aza-dC, suggesting that the mitotic maintenance of differences in AI between clones in such loci was due to some mechanism other than DNA methylation.

Comparison of AI changes of the same genes across clones revealed a striking property. Depending on the initial AI, allelic bias in a given clone could shift towards balanced expression (AI=0.5) or towards stronger bias. However, when considered together, AI values converged across clones after exposure to 5-aza-dC (**Fig.4F**). Principal component analysis for all genes with AI shifts showed that transcriptome-wide AI states of clones became closer to each other after demethylation (**Suppl. Fig.S16**). The target AI to which clones converged varied between genes and encompassed the whole range of AI values, including extreme allelic bias for some genes (inferred target values shown as green boxes in **Fig.5** and **Suppl. Fig.S17**). In clones where an MAE gene was already in the target AI state (e.g., *Col6a5* in Abl.3 and Abl.4, or *Casp6* in Abl.1 and Abl.4; **Fig.5**), there were no further AI shifts after DNA demethylation.

**Figure 5.**
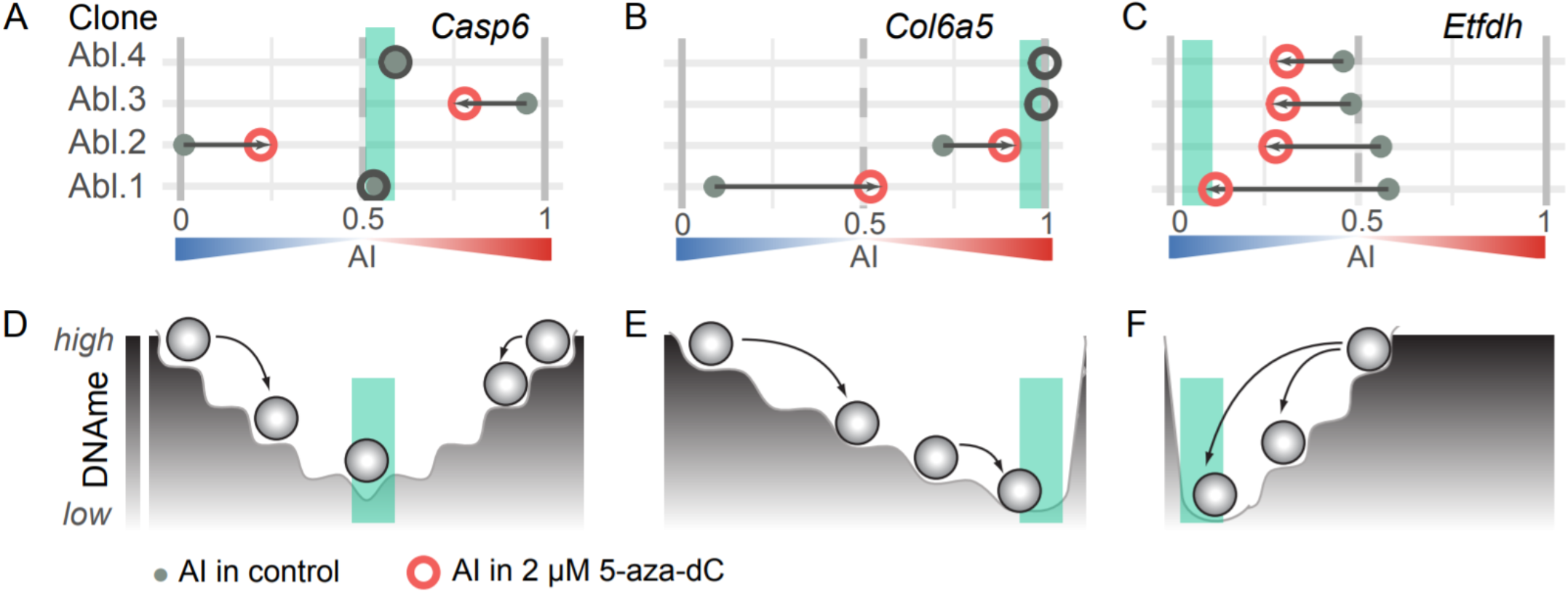
DNA methylation as a rheostatic mechanism of the mitotic maintenance of allele-specific expression. **(A-C)** Examples of genes with significant changes in AI across four lymphoid clonal lines. All genes with statistically significant shifts (after correction for AI overdispersion) are shown in **Suppl. Fig.S17.** AI values in control *(filled grey circles*) and after exposure (*open circles*) are shown (no shifts are in grey). Same Abl.1 data as in **Fig.4**. Clones Abl.2, Abl.3 and Abl.4 were exposed to 0.2 μM 5-aza-dC. *Green box* denotes imputed common end states. **(D-F)** Diagram of interaction between cis-regulatory landscape and DNA methylation (DNAme) which determines the rheostatic control of allele-specific expression. There are multiple stable DNAme states, and corresponding AI in expression. Decreases in DNAme lead (arrows) to convergence of states between clones, with complete loss of DNAme leading to common AI state (*green box*).

These observations are consistent with a simple model in which the AI for many MAE genes are set at different values across clones and are maintained via DNA methylation of regulatory sequences in *cis* to the affected genes. Demethylation causes these AI values to converge to a genetically determined “target” state, with partial demethylation resulting in an AI value between the start and the target states (**Fig.5**).

## DISCUSSION

Using a screening-by-sequencing approach, we identified DNA methylation as a key mechanism involved in the mitotic maintenance of monoallelic expression in clonal cell lineages of mammalian cells. Dnmt1-dependent maintenance DNA methylation offers a straightforward explanation for MAE stability, since it is a very stable form of molecular memory: cytosine methylation in the *Cryptococcus* genome has apparently been maintained for millions of years in the absence of *de novo* methylation (35). We propose a simple model (**Fig.5**), with the allele-specific regulatory landscape defined by genetic variation (possibly in interaction with epigenetic mechanisms), while a specific state of a clonal cell population depends on DNA methylation. Note that this model predicts specific regulatory elements located in *cis* to the affected genes. Such genomic elements would offer a simple explanation to the evolutionary conservation of MAE status of genes across human populations (18) and between human and mouse (12, 13, 15). When and how DNA methylation is established in these regulatory regions remain to be uncovered.

Not all assessed MAE genes were affected by DNA demethylation, suggesting that MAE maintenance for some loci involves other mechanisms instead of (or in addition to) DNA methylation. This offers one likely explanation of the previous observations that DNA demethylating agents did not affect allelic imbalance in any of the several assessed MAE genes (8, 9). Consistent with the idea of additional mechanisms of MAE maintenance, a SIRT1 activator BML-278 and a sirtuin inhibitor, salermide, appeared as primary hits in our screen (see **Suppl. Fig.S3**), suggesting that expanded application of the Screen-seq strategy can uncover such additional mechanisms.

We assessed the genome-wide impact of DNA demethylation on the allelic imbalance in the transcriptome of clonal lymphoid cells. Using a new, highly stringent approach to allele-specific RNA-seq analysis (22), we found AI shifts in over 600 autosomal genes between four analyzed clones. Interestingly, only six X-linked genes showed small but statistically significant changes (**Suppl. Fig.S18**), suggesting that DNA methylation-dependent mitotic maintenance of AI is easier affected in autosomal genes than X-chromosome inactivation.

Significant impact of DNA demethylation drugs on allele-specific expression in lymphocytes is of particular importance since both 5-aza-2’-dC and 5-azaC are used in the clinic to treat acute leukemia and other malignancies (28). Notably, concentrations of these compounds in our experiments (0.2 – 1.0 μM; **Fig.3**) are similar to that measured in the patients’ plasma [5-aza-dC at ~60 ng/ml, about 0.25 μM (36)]. Our findings thus imply that DNMT inhibitors likely affect gene regulation in patients in ways that would be undetectable without allele-specific analysis. Similarly, we note that large shifts in AI were often independent of changes in overall RNA abundance (see **Suppl. Fig.S10, Suppl. Fig.S12** and **Suppl. Table S8**). This suggests that analyses of allele-specific gene regulation in polyclonal and monoclonal cell populations should lead to new translational insights.

Stringent quantitative analysis of RNA-seq data reveals a more complex landscape of mitotically stable clonal diversity in allele-specific gene regulation than is implied by monoallelic/biallelic dichotomy. This complexity suggests that the biologically relevant questions concern molecular mechanisms and functional impact on cellular function, rather than arbitrary thresholds. In different clones, AI of a gene can span a range of values, with epigenetic regulation acting as a rheostat rather than an on/off switch. This extends the idea of the rheostatic role of DNA methylation, proposed for the regulation of the overall abundance of transcripts (37, 38).

## METHODS

### Cell culture

v-Abl pro-B clonal cell lines Abl.1, Abl.2, Abl.3 and Abl.4 (13) were cultured in Roswell Park Memorial Institute medium (Gibco), containing 15% FBS (Sigma), 1X L-Glutamine (Gibco), 1X Penicillin/Streptomycin (Gibco) and 0.1% β-mercaptoethanol (Sigma).

### Drug treatment

The SCREEN-WELL® Epigenetics arrayed drug library was purchased from Enzo Life sciences (BML-2836). The Abelson clone Abl.1 was treated with the entire drug library (**Suppl. Table S3**) at concentrations of 1 μM, 10 μM and 20 μM in order to encompass a wide enough range of concentrations where the drugs are potentially pharmacologically active. Cultures were treated for 21 days where media was changed every second day. After following up hits from the initial drug screen, 5-aza-2’-deoxycytidine (5-aza-dC, Sigma, A3656) was diluted in DMSO at a concentration of 10mM and Abl.1 cells were treated using a concentration range of 10 nM to 20 μM 5-aza-2’-deoxycytidine. Cells were treated for a total of 21 days where media was changed every 2 days and samples of ~1×10^5^ cells were harvested for RNA extractions on days 2, 5, 7, 9, 12 and 14. Viable cells were counted using trypan blue solution (Gibco™) on Countess™ II FL Automated Cell Counter machine (Life Technologies). For all treatments, drugs were solubilized in DMSO and dilutions were made to ensure the final DMSO added to cultures was 1% (v/v). For exposure/recovery experiment (**Fig.3**), 5-aza-dC was dissolved in water.

### RNA and DNA preparation

For all Abelson monoclonal cultures, RNA was extracted from cells using a magnetic bead-based protocol using Sera-Mag SpeedBeads™ (GE Healthcare). Isolated RNA was DNase-treated with RQ1 DNase (Promega). First strand cDNA synthesis was done using Episcript™ RNase H-reverse transcriptase (Epicentre) where RNA samples were primed with random hexamers (NEB). Both DNase treatment and cDNA synthesis were performed using manufacturer specifications with minimal modifications. For RNA preparation from mouse spleen, cells were extracted by crushing the whole spleen using the back of 1 ml syringe plunger in 40 μM nylon filter and washing the strainer with 1X PBS (Phosphate-buffered saline, Sigma) to collect cells. Cells from spleen were spun down and RNA was extracted using Trizol reagent (Invitrogen). Genomic DNA extractions for testing the sensitivity of Screen-seq were performed using the salting out method (39) and for reduced representation bisulfite sequencing (RRBS) were performed using Sigma GenElute kit (G1N10-1KT). RT-qPCRs were performed using iTaq™ Universal SYBR® Green Supermix (BioRad) using manufacturer’s protocol on a 7900HT Fast Real-Time PCR system (Applied Biosystems Inc.). All primers used in this study were ordered from Integrated DNA Technologies and their sequences are listed in **Suppl. Table S2**.

### Screen-seq methodology

A targeted sequencing method similar to that described in Nag et al (2013), was used to assay multiple genes simultaneously for assessing allele-specific expression. Here, we assayed 23 genes. The assay involved RNA extraction, cDNA synthesis (**Fig.1B**), two rounds of PCR amplification and Illumina sequencing. After magnetic bead-based RNA purification, cDNA synthesis was performed within each well of a 96-well plate, separately using EpiScript™ Reverse Transcriptase (EpiCentre Biotechnologies) using both random hexamers (NEB) and UMI-tagged oligo-dT primer with universal tail (**Suppl. Table S2**) using manufacturer’s instructions. Half the portion cDNA products were transferred to a separate 96-well plate. Gene-specific multiplex PCR are performed in both the plates using Phusion U multiplex Master Mix (ThermoFisher, F562L, Waltham, MA). Two types of multiplexed readouts were generated within each plate: readouts without UMI and readouts with 3’-UMI. For the multiplex readouts without UMI, target genes that contain the SNP(s) differentiating the maternal and paternal allele, were amplified using gene-specific primer pairs containing one of two universal tails (UT1 or UT2, **Suppl. Table S2**). For the multiplex readouts with 3’-UMI, the forward primers were gene-specific and contained universal tail UT2 (**Suppl. Table S2**). They were always positioned near the SNP of interest. Reverse primer for these genes were complimentary to the universal tail UT1. These readouts were always constrained to the 3’ end of the transcript. These two types of multiplex readouts were not generated for all readout genes. A list the readout genes for which the multiplex assay was used is given in **Supp. Table S1**. MPprimer primer design program (40) was used to design the non-UMI multiplex PCR assay. We computationally generated an input form that would 1) constrain our SNP(s) of interest within 135 base pairs from one end of the amplicon, 2) mask repetitive regions, 3) prevent the design of primers pairs that exist within more than one exon and 4) ensure that the total fragment size for each readout falls within 250-500 base pairs. Once the gene-specific primer sequences were designed, the universal tails were added. Primers generated were tested for specificity and primer dimerization using MFEprimer (41) and also experimentally validated. The two groups of multiplex products from the gene-specific PCR were combined and carried over as templates to the second PCR which is performed using Phusion® High-Fidelity DNA Polymerase (New England Biolabs Inc., M0530L, Ipswich, MA) that barcodes each well/perturbation separately. These reactions use primers that target the universal tails (UT1 and UT2) of the readouts amplified in the first multiplex PCR and add a six-nucleotide barcode, a seven-nucleotide spacer and an Illumina primer dock (**Suppl. Table S2**). Combinatorial barcoding was achieved by using a pair of unique forward and reverse primers, which tag each sample with a unique barcode combination. These barcode combinations allowed pooling of samples in the subsequent steps of the assay. Once pooled, the readout library was cleaned up using magnetic beads at a bead to sample ratio of 1.2 to get rid of primer dimer bands <150bp in size. The sample was then carried over as a template into a third PCR reaction which adds Illumina adapters.

We observed high accuracy of multiplexing and barcoding steps of Screen-seq by comparing the AI calculated from Screen-seq and expected AI for a range of pure 129 and Cast parental genomic DNA mixes for all genes (**Suppl. Fig.S1**). A good correlation was observed between the reads with UMI and without UMI for readout genes tested using both methods. For this, Screen-seq was performed for a range of RNA mixes from pure 129 and Cast mice spleen. Allelic imbalance (AI) calculated from Screen-seq for *Adamtsl4* and *Adnp2* showed good correlation with the expected AI, and also between UMI and non-UMI assays (**Suppl. Fig.S2**). *Smtnl2* and *Dnajc12* showed low expression in mice spleen tissue and hence comparison could not be made. Finally, the assays for genes we selected had to combine successfully in multiplexed PCR.

### Screen-seq data analysis

After Screen-seq libraries were prepared as described above, they were sequenced at the UMass Boston and Center for Cancer Systems Biology (CCSB) sequencing core on Illumina HiSeq 2500 and MiSeq, respectively using four-color reagent kits. From the P7 adapter end, 65nt were sequenced (Read 1), including one of the two barcodes for encoding plate wells (and the UMI, where appropriate). From the P5 adapter the remaining 135nt were sequenced (Read 2), covering the second well-encoding barcode and the cDNA amplicon containing the interrogated SNP. In addition, standard Illumina barcodes were used to distinguish individual plates within the overall pooled library, with demultiplexing before further processing. Reads were aligned using *bowtie2* (42) against mm10 mouse genome assembly. The resulting BAM files were processed using custom Perl scripts to extract allele-specific, UMI-corrected counts for each gene and each well.

To identify primary hits (outliers in **Figs.1E and 1F**), the allele-specific counts were analyzed using custom R scripts. Briefly, for each gene, point AI estimates for all drug conditions were considered together to determine median AI and the interquartile range (IQR = Q3 − Q1, with Q1 and Q3 the 25th and 75th percentiles). Observations with counts under 50 were filtered out (an observation consists of allelic counts for one gene in one well). A common practice for identification of outliers is to use values below Q1−1.5×IQR or above Q3+1.5×IQR (44). We used a more stringent threshold of 3×IQR, to reduce the likelihood of false positive hits. Complete results can be found in **Suppl. Table S5**.

### Droplet Digital PCR

Droplet digital PCRs (ddPCRs) were performed on QX200 ddPCR system (BioRad) for absolute quantification of 129 and Cast alleles using manufacturer-recommended settings. C1000 Touch™ thermal cycler was used to perform amplification within droplets. SNP-specific TaqMan assays (IDT; sequences in **Suppl. Table S2**) were designed manually. We first validated all TaqMan assays experimentally using homozygous Cast and 129 cDNA and optimized reaction conditions for each assay using Abl.1 clonal cell line cDNA, including Tm of each primer-probe mix by performing thermal gradient PCR. Finally, we tested the specificity of this method by using known quantities of left kidney cDNA from homozygous 129 and Cast mice parents and comparing it with the estimated allelic imbalance from ddPCR. To determine the false-positives, we made 2-fold dilutions of these samples starting from 1 ng cDNA till its 1/16^th^ dilution. Results demonstrated our ability to precisely measure allelic imbalance in samples with 30 copies/μl using ddPCR. cDNA was prepared from around 100,000 cells and 8μl template cDNA (1/4^th^ of eluted sample) was used per reaction. Gating for clusters with maternal and paternal alleles was decided by comparing the fluorescence intensity individually for the maternal and paternal probes in homozygous 129 and Cast tissue samples. Data was processed using QuantaSoft v.1.6 (Bio-Rad). Inverse fractional abundance given displayed by the Quantasoft software was divided by 100 and noted as AI measurement [mat/(mat+pat)] from ddPCR.

### shRNA infection

Two shRNA vectors targeting *Dnmt1* (SHR000038801.1_TRC001.1 and SHR000373188.1_TRC005.1) and a control empty vector (NUL003.3_TRC021.1) packaged in lentiviral vectors obtained from the Genetic Perturbation Platform at the Broad Institute were tested. The optimal multiplicity of infection (MOI) was determined by infecting Abl.1 cells with pLKO_TRC060 lentiviral vector expressing eGFP. Abl.1 cells were infected with 3 shRNA vectors (2 targeting Dnmt1 and 1 control) individually on day 1 at the optimal MOI under normal growth conditions in the presence of 8 μg/ml polybrene and spun at 800×g for 90 minutes at 37°C. The next day the media was changed and media containing 2 μg/ml of puromycin was added on day 2. Selection was maintained continuously afterwards, and media changes were done every 2-3 days. Cells were harvested on day 12 and 19 after infection, and RNA was extracted.

### Estimation of allelic imbalance in RNA-seq, RRBS and exome sequencing

5×10^6^ cells treated with concentrations of 0.2 μM, 2 μM and 10 μM 5-aza-2’-deoxycytidine were harvested on days 1, 2 and 5. Live cells were separated from debris by sucrose gradient centrifugation (Histopaque®-1077, Sigma). RNA was extracted from 2×10^5^ live cells, and the remaining live cells were washed with 1X PBS and flash frozen on dry ice for genomic DNA extractions.

Libraries for RNA-seq were prepared for cells collected on day 2, using at least two technical replicates from the same RNA (5 replicates for Abl.1 cells treated with 2 μM 5-aza-dC), using SMARTseqv4 kit (Clonetech), starting with 10 ng input RNA for each library according to manufacturer’s instructions. Library preparation, QC and sequencing were performed at the Molecular Biology Core Facilities at Dana-Farber Cancer Institute. Single-end 75bp reads were generated using a Nextseq 500 instrument (Illumina).

Allele-specific gene expression analysis was performed using ASEReadCounter* and Qllelic version v0.3.1 pipeline described in (22). Briefly, RNA-seq reads were aligned with STAR aligner v.2.5.4a using imputed parental genomes as reference, with default quality filtering. Only uniquely aligned reads were used. Allele-specific coverage over SNPs was counted using *samtools mpileup* and further processed using ASEReadCounter* tool based on the GATK pipeline. All exons belonging to the same gene were merged into a single gene model based on RefSeq GTF files (GRCm38.68 and GRCh37.63); overlapping regions that belong to multiple genes were excluded. AI point estimate per gene obtained as a proportion of maternal gene counts to total allelic gene counts. Differences in AI were accepted as significant after accounting for experiment-specific overdispersion, estimated using Qllelic v0.3.1 analysis of replicate libraries. Complete results can be found in **Suppl. Table S9**.

To control for the possible loss of heterozygosity, exome sequencing was performed on genomic DNA for all clones. Library preparation, QC and sequencing (50x) were performed at LC Sciences (TX, USA). Exome capture was performed using SureSelect (Agilent Technologies) following the vendor’s recommended protocol. Paired-end 150bp reads were generated using a Hiseq X Ten sequencing instrument (Illumina). Genes with total allelic counts of <10 and those with nominal AI ≥0.7 or ≤0.3 were excluded from respective clones before comparing RNA-seq data between clones.

For Reduced Representation Bisulfite-seq (RRBS), libraries were generated from 50 ng input genomic DNA using a scaled-down (half reactions) of the NuGEN Ovation RRBS Methyl-Seq System (Tecan) following the manufacturer’s recommendation. Libraries were PCR amplified for 11 cycles. Paired-end 100bp reads were generated using HiSeq 2500 instrument (Illumina). Reads were aligned to the mouse mm10 genome using BSmap3 with flags -v 0.05 -s 16 -w 100 -S 1 -p 8 -u. Custom scripts written in Perl were used to calculate the methylation percentage for CpGs covered by 4 or more reads (43) at locations of known SNPs. Briefly, VCF files containing SNPs between 129 and Cast strains were filtered to exclude calls that did not pass minimum requirements as well as C→T or G→A calls. For each SNP, RRBS reads that overlapped that SNP were extracted, and the methylation status of genomic cytosines was calculated by dividing the number of unconverted (methylated) cytosines (C) by the total number of unconverted (C) or converted (T) cytosines. The methylation status of all cytosines on reads overlapping a SNP were aggregated by SNP status to create a methylation average for the reference and alternate genotype. Complete results can be found in **Suppl. Table S10**.

## Supporting information

Main Supplemental File

## Data availability

RNAseq and RRBS data is deposited at NCBI Gene Expression Omnibus (GEO) repository under accession number GSE144007 (subseries: GSE144005 for RNAseq and GSE144006 for RRBS).

## ACKNOWLEDGMENTS

We thank Alex Bortvin, Arun Chavan, Mary Gehring, David Hill, Mitzi Kuroda and members of the Gimelbrant lab for valuable comments, Matt Warman for generously sharing a ddPCR system, and Kim Blanchard and Arman Mohammed for technical help.

This work has been supported by NIH grants R21HD081675 and R01GM114864 to AAG and Marie Curie fellowship (EU project 752806) to CFAP.

## Notes

### Competing Interest Statement

The authors have declared no competing interest.

