## Supplementary material for "DNA methylation is a key mechanism for maintaining monoallelic expression on autosomes": Main Supplemental File

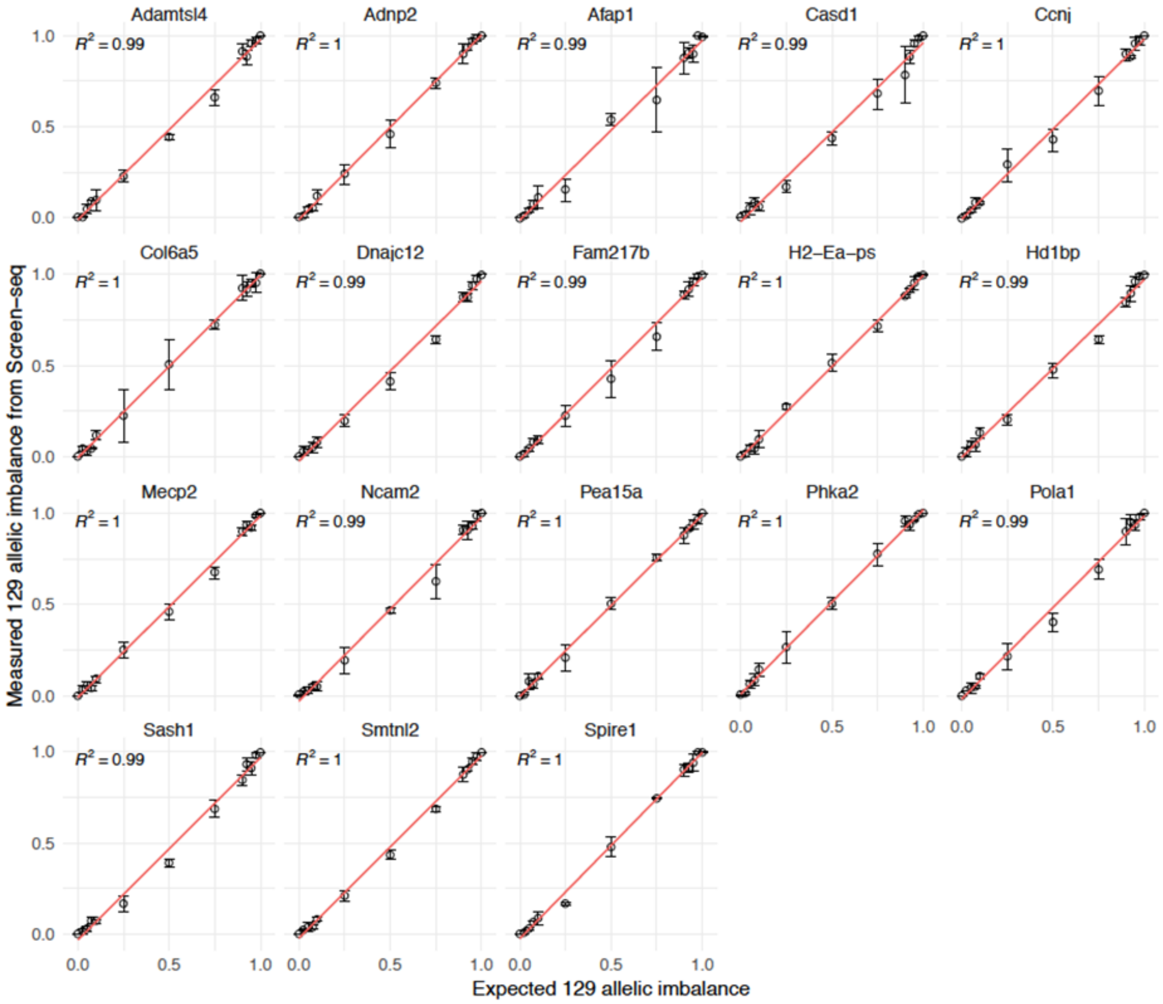

**Supplemental Figure S1. Expected vs measured allelic imbalance (AI) from Screen-seq. Related to Figure 1B.** Since no perturbations are known that can change AI in any locus, much less in all targeted loci, for the control experiments we titrated known mixes of genomic DNA from liver tissue of the parental mouse strains, 129S1/SvImJ and Cast/EiJ. Allelic imbalance was measured as AI= [129 counts]/[129 + Cast counts]. gDNA mixes ranging from 0 to 0.1 and 0.9 to 1 in 0.025 increments, as well as 0.25, 0.5, and 0.75, were made from parental liver tissue and shown on X-axis (Expected AI). For AI measured from Screen-seq (Measured AI, Y-axis), data points are an average of 3 biological replicates and error bars show standard deviation. *Red* line denotes linear fit to smoothen all data points.

Expected and measured AI were highly concordant ($R^2 \geq 0.99$) at >1000 reads/SNP for readout genes.

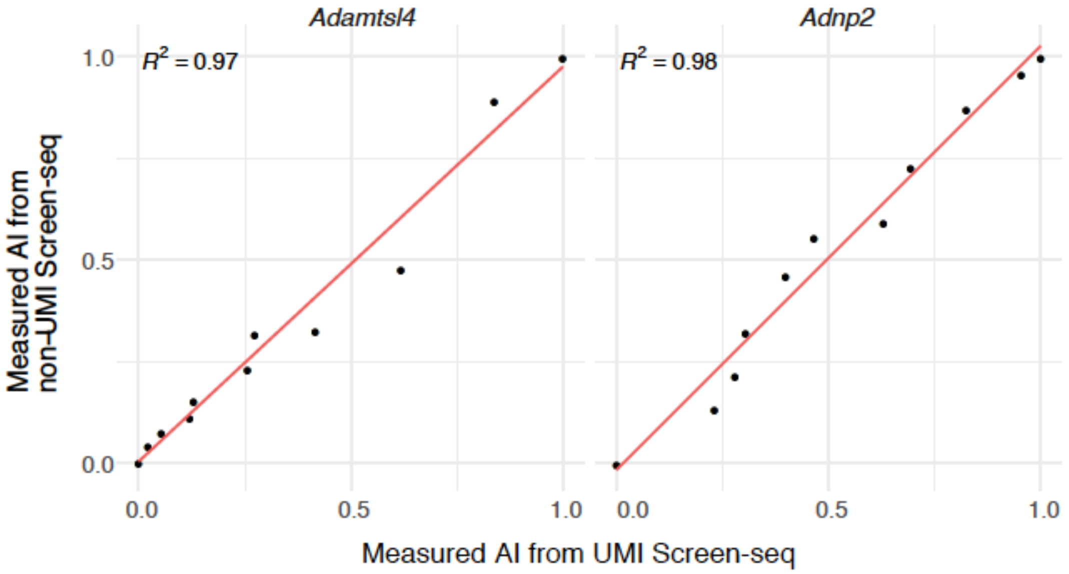

**Supplemental Figure S2. UMI vs non-UMI based AI measurement from Screen-seq. Related to Figure 1B.** We compared AI sensitivity for UMI and non-UMI assays, by designing both types of assays for a subset of genes where the position of SNPs allowed that. For this, we used mixes prepared from total RNA from the spleens of the mice of the parental mouse strains. Both types of assays were designed for a subset of genes (*Adamtsl4, Adnp2, Dnajc12, Smtnl2*) where the position of SNPs allowed that. AI measurements were highly concordant between the UMI and non-UMI assays ($R^2 \geq 0.97$) at >1000 reads/SNP.

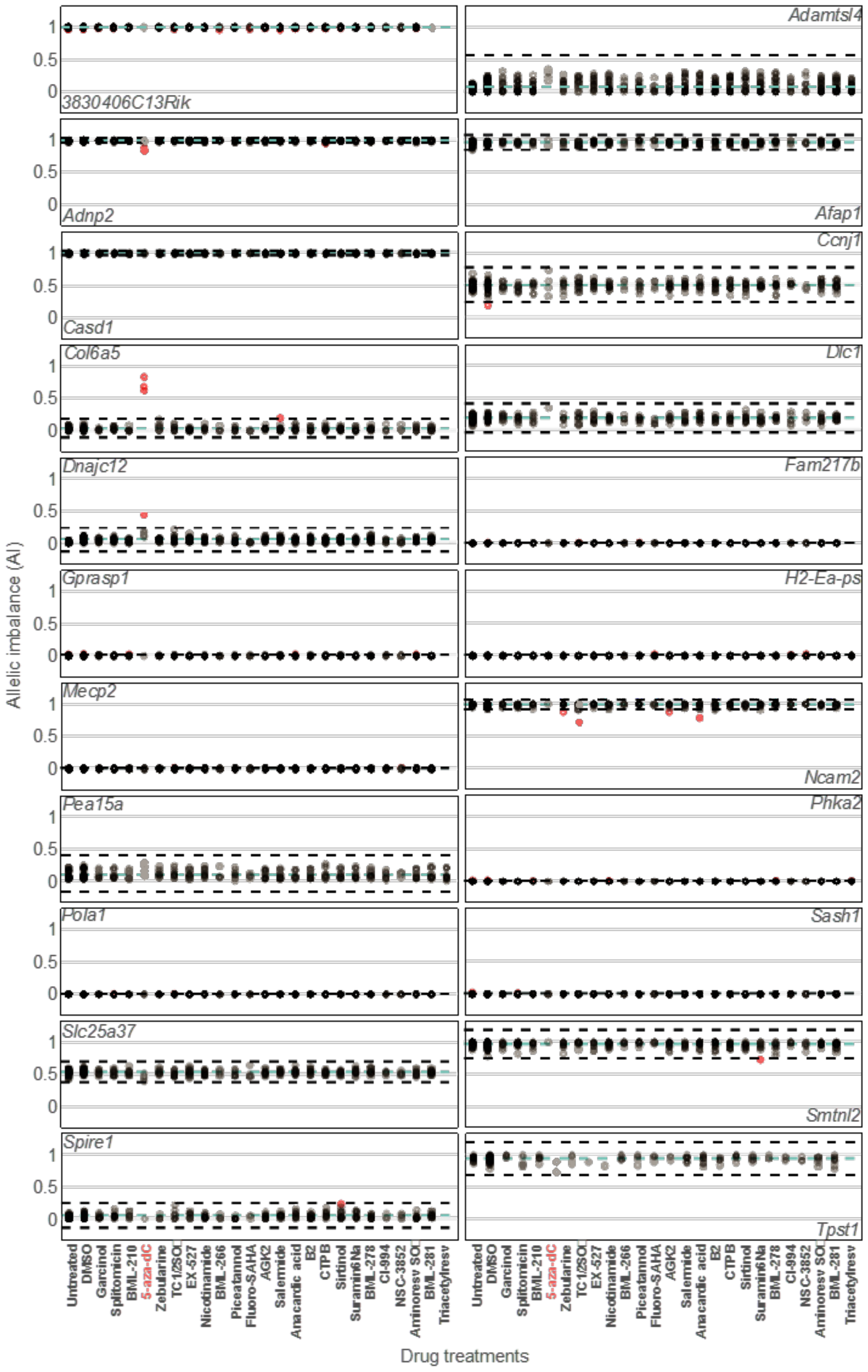

**Supplemental Figure S3. Screen-seq results for 23 readout genes. Related to Figure 1E.**
All time and concentration points for a single drug shown in the same column. Each point shows AI in one condition. *Blue line*: mean AI for a gene across all conditions; *black dashed lines*: [Q1-3×IQR] and [Q1+3×IQR] (inter-quartile range); *red* points: outlier AI values (hits). See **Suppl. Table S4** for drug hits and **Suppl. Table S5** for complete Screen-seq results.

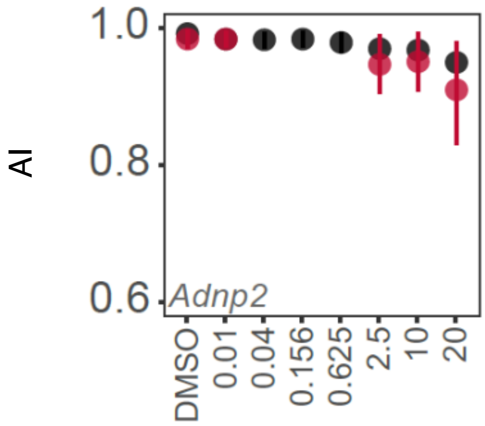

**Supplemental Figure S4. *Adnp2* allele imbalance in Abl.1 cells on 5-aza-dC treatment with ddPCR. Related to Figure 2D.**
Biological replicate analysis with 5-aza-dC in Abl.1 cells after 2- and 5-days exposure (*black* and *red* points, respectively) using ddPCR. Summary of ddPCR analysis for *Adnp2,* another MAE readout gene that showed up as a hit in Screen-seq on treatment with 5-aza-dC. Concentrations of 5-aza-dC (in µM) are shown on X-axis.

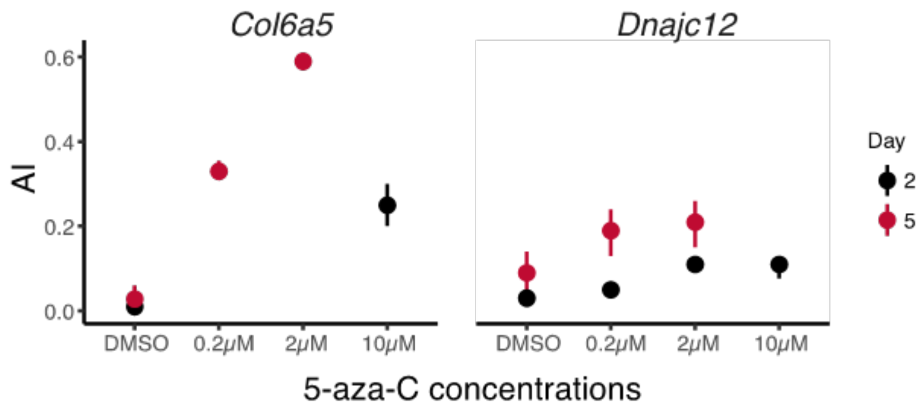

**Supplemental Figure S5. Effect of 5-aza-cytidine (5-aza-C) on allelic imbalance of *Col6a5* and *Dnajc12* measured with ddPCR. Related to Figure 2D.**
Effect of 5-aza-cytidine (5-aza-C), a closely related analog of 5-aza-dC, on AI. Summary of ddPCR analysis in Abl.1 cells for *Col6a5* is shown on *left* and for *Dnajc12* is shown on *right*.

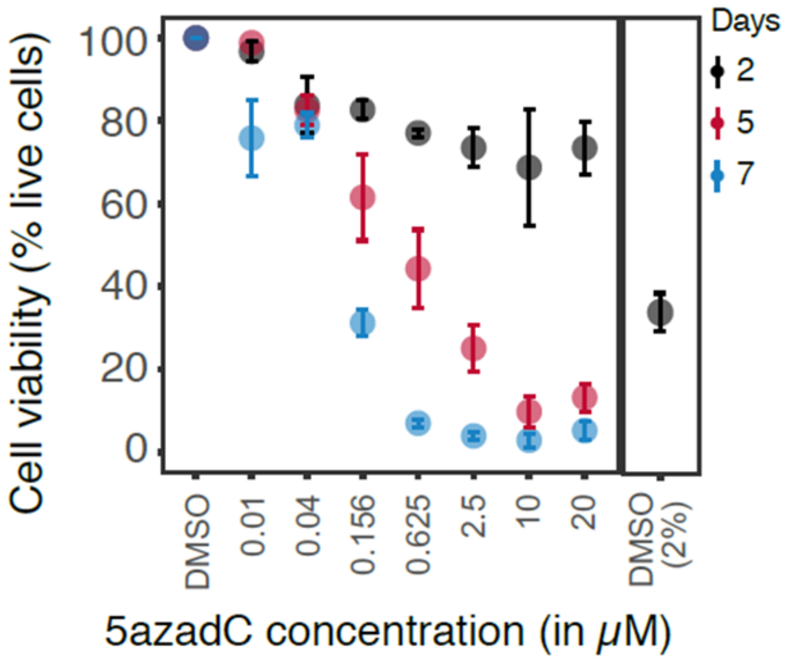

**Supplemental Figure S6. Effect of 2% DMSO on cell viability of Abl.1 cells. Related to Figure 2D.**
Effect of 5-aza-dC and 2% DMSO on cell viability of Abl.1 cells. 1% DMSO was used as the drug solvent. Data points are an average 3 biological replicates and error bars show S.E.M (standard error of mean).

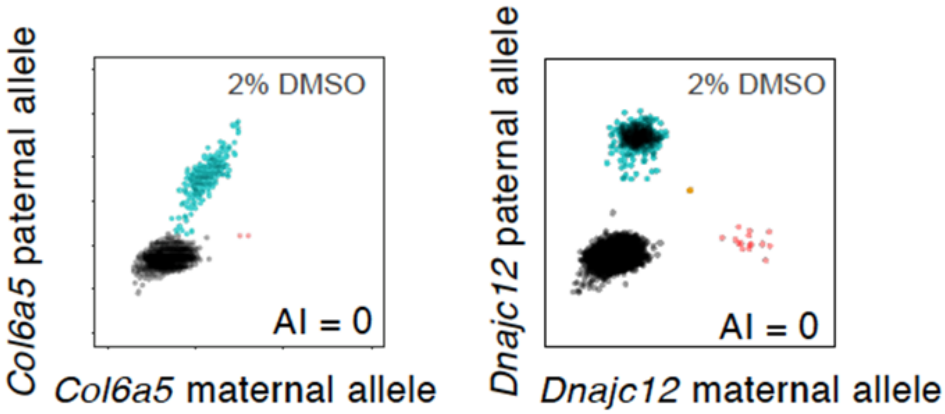

**Supplemental Figure S7. Effect of high concentration of DMSO (2%) on allelic imbalance of *Col6a5* and *Dnajc12* measured with ddPCR. Related to Figure 2D.**
ddPCR scatterplots for 20,000 droplets targeting readout gene *Col6a5* (*left*) and Dnajc12 (*right*) when Abl.1 cells were treated with 2% DMSO (*right*) in growth medium for 2 days. *Black*: empty droplets; *blue*: droplets with Cast allele amplified; *red*: droplets with 129 allele amplified. Ratio of red:red+blue droplets shown as AI.

**(A)**

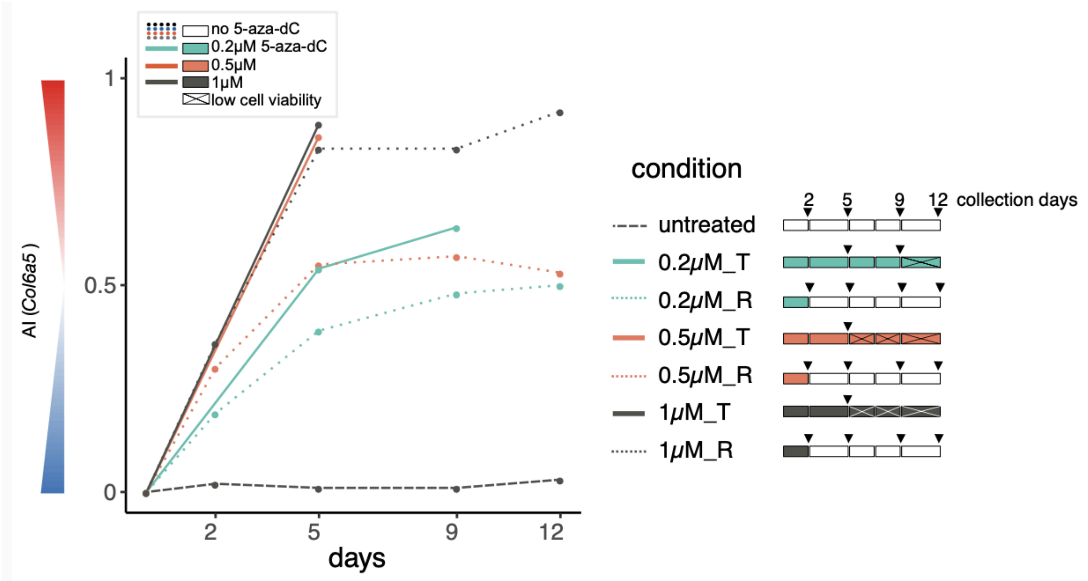

**(B)**

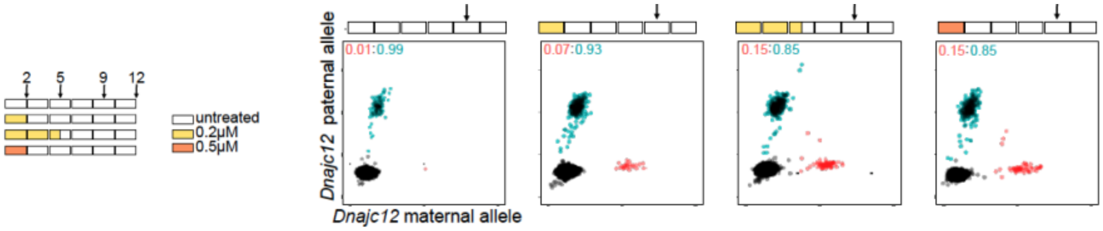

**(C)**

| | with drug | | | | d2 recovery | | | | d5 recovery | | | |
|---|---|---|---|---|---|---|---|---|---|---|---|---|
| days | 0 | 0.2 | 0.5 | 1 | 0 | 0.2 | 0.5 | 1 | 0 | 0.2 | 0.5 | 1 |
| 1 | 94% | 84% | 80% | 76% | | | | | | | | |
| 2 | 92% | 67% | 64% | 69% | | | | | | | | |
| 3 | 89% | 49% | 33% | 21% | 68% | 45% | 9% | 26% | | | | |
| 4 | 69% | 37% | 7% | 27% | 65% | 46% | 22% | 6% | | | | |
| 5 | 69% | 38% | 12% | 7% | 68% | 72% | 25% | 9% | | | | |
| 6 | 71% | 30% | 5% | 0% | 43% | 72% | 24% | 3% | 74% | 31% | 0% | 0% |
| 7 | 80% | 11% | 0% | 2% | 78% | 86% | 40% | 7% | 79% | 48% | 0% | 5% |
| 9 | | | | | 86% | 82% | 68% | 46% | 90% | 58% | 0% | |
| 12 | | | | | 74% | 66% | 75% | 58% | 75% | 77% | 0% | |

**Supplemental Figure S8. 5-aza-dC treatment-and-recovery experiment in Abl.1 cells - Complete results. Related to Figure 3.**

(A) *Col6a5* gene. Drug treatment setup is shown next to the line plots. Cells were exposed to 0.2, 0.5 or 1 μM 5-aza-dC (denoted by colors) in growth medium. Regular media changes were done as shown by breaks in the boxes in drug treatment setup. Cells treated with 5-aza-dC continuously are shown as solid colored lines (condition labeled as *5-aza-dC concentation*_T). AI in cells that were moved to growth media without any drug after drug exposure of 2 days are shown as dotted lines (condition labeled as *5-aza-dC concentation*_R). AI measurements were performed with ddPCR.

(B) *Dnajc12* gene. Same setup as (A). Mitotic memory of Dnajc12 AI was observed after 5-aza-dC exposure and recovery on day 9. Scatterplots showing Dnajc12 AI in Abl.1 cells were generated using ddPCR.

(C) Abl.1 cell viability during the complete experiment. d2 recovery implies cells were exposed to drug for 2 days, after they were washed twice and transferred to growth medium containing no drug.

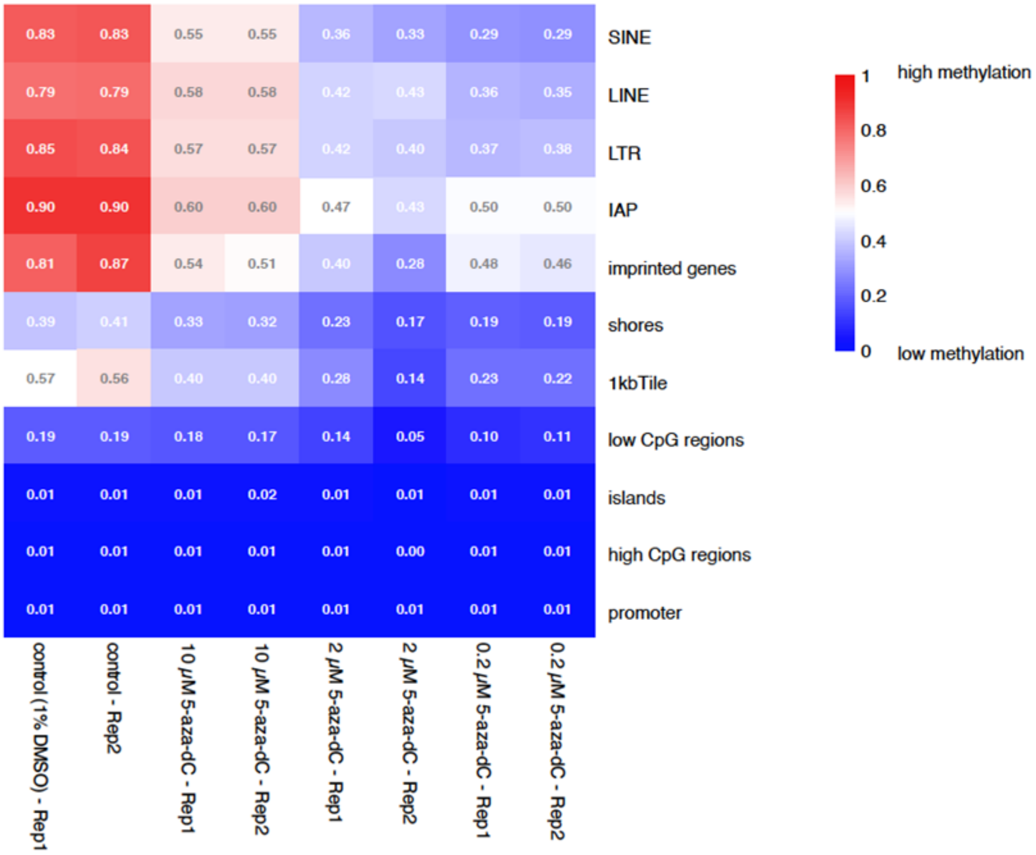

**Supplemental Figure S9. Genome-wide DNA methylation in Abl.1 cells with RRBS. Related to Figure 4D.** Median methylation values for genomic features, clustered by sample (x-axis) and genomic feature (y-axis) using hierarchical clustering. Genomic features include repetitive elements (SINE, LINE, LTR, IAP (2)), curated imprinting control regions, CPG Islands (1) and CPG Shores (2kb surrounding CpG Islands), 1kb tiles across the genome, high- and low-CPG promoters, and all promoters. The methylation value for each genomic region was calculated as the average of methylation values of CpGs in that region. CpGs were required to be covered at 3x in at least 11/14 samples to contribute to the region average. Note that high 5-aza-dC concentration of 10µM leads to lesser demethylation than 2µM 5-aza-dC, likely due to higher toxicity of 10µM 5-aza-dC (**see Suppl. Fig.S6**).

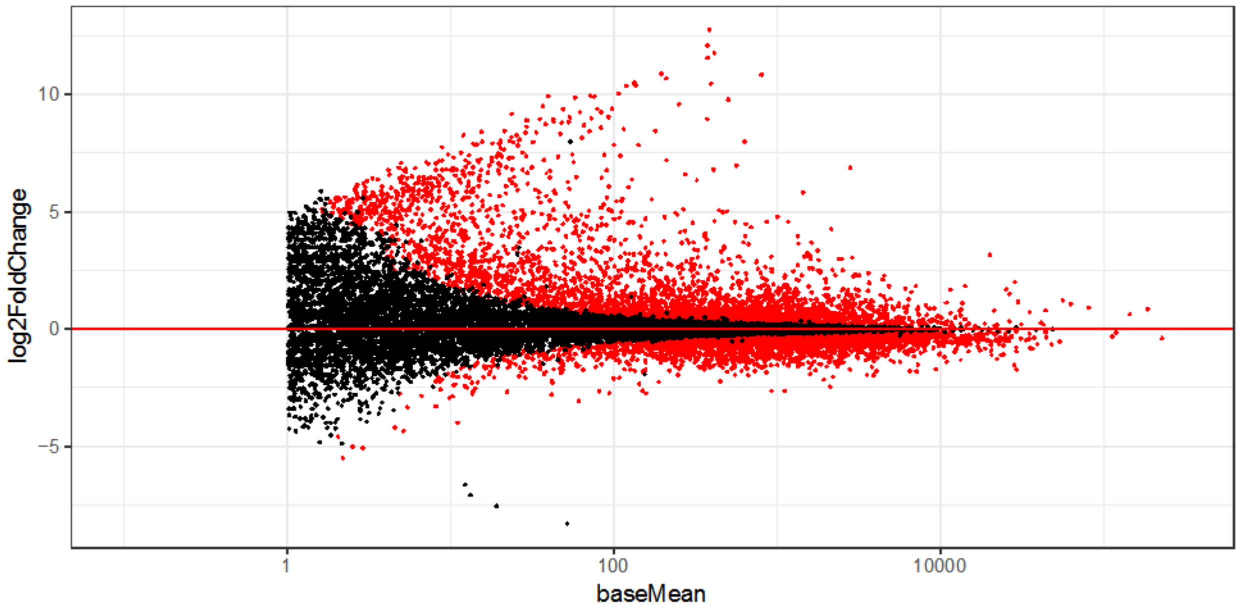

**Supplemental Figure S10. Genome-wide RNA abundance in Abl.1 cells on 5-aza-dC treatment with RNA-seq. Related to Figure 4E.**
MA plot showing increase in RNA abundance on 10μM 5-aza-dC treatment in Abl.1 cells. Each point is a gene. Y-axis shows $log_2$ fold change in gene expression between control (1% DMSO) and 10μM 5-aza-dC conditions. X-axis shows gene expression (TPM values) in control condition. Genes that changed expression ($P_{adj} < 0.05$) are shown in red.

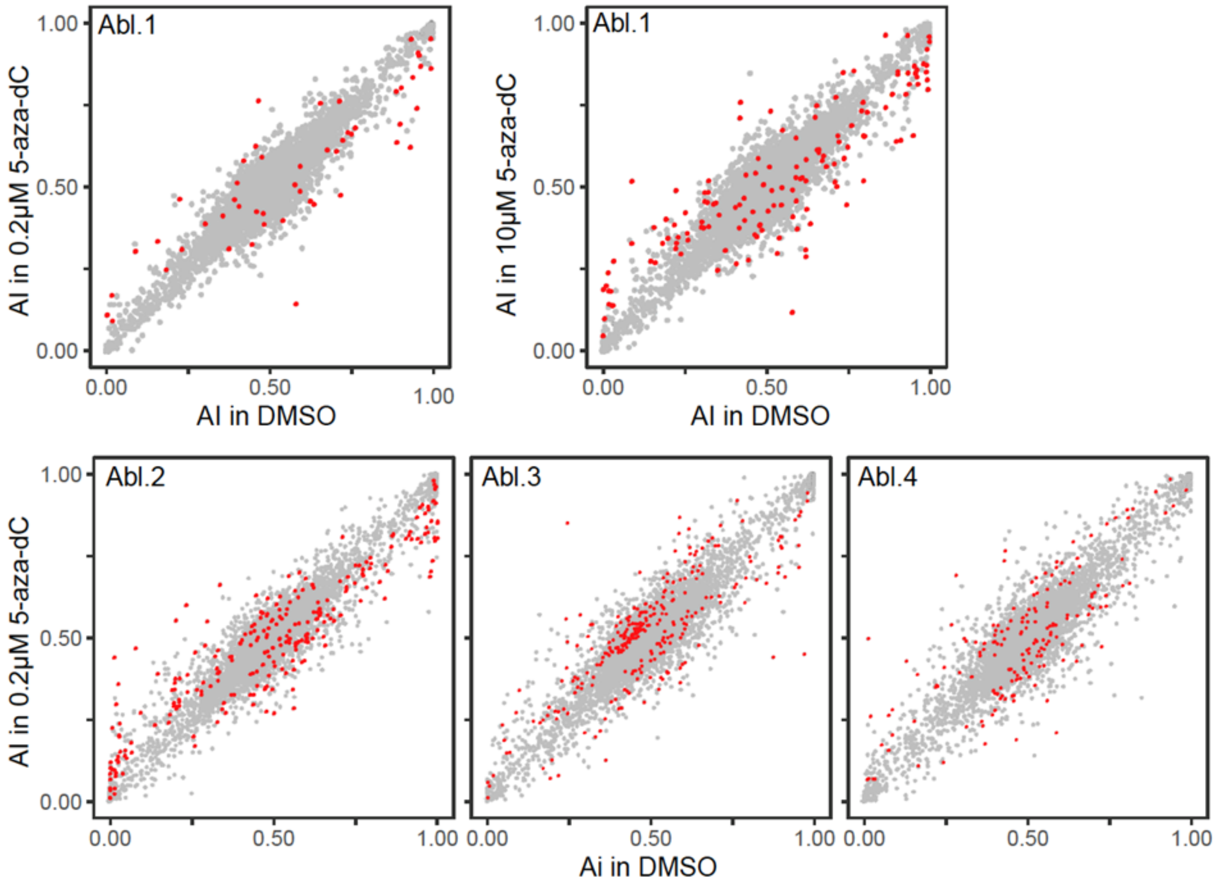

**Supplemental Figure S11. Genome-wide allele-specific RNA expression of all clones in control and 5-aza-dC treatment with RNA-seq. Related to Figure 4A and 4F.**
AI values for gene expression in Abl.1, Abl.2, Abl.3 and Abl.4 cells in control (horizonal axis) and 5-aza-dC (concentration mentioned on vertical axis). Genes passing stringent significance threshold for differential AI are shown in *red*.

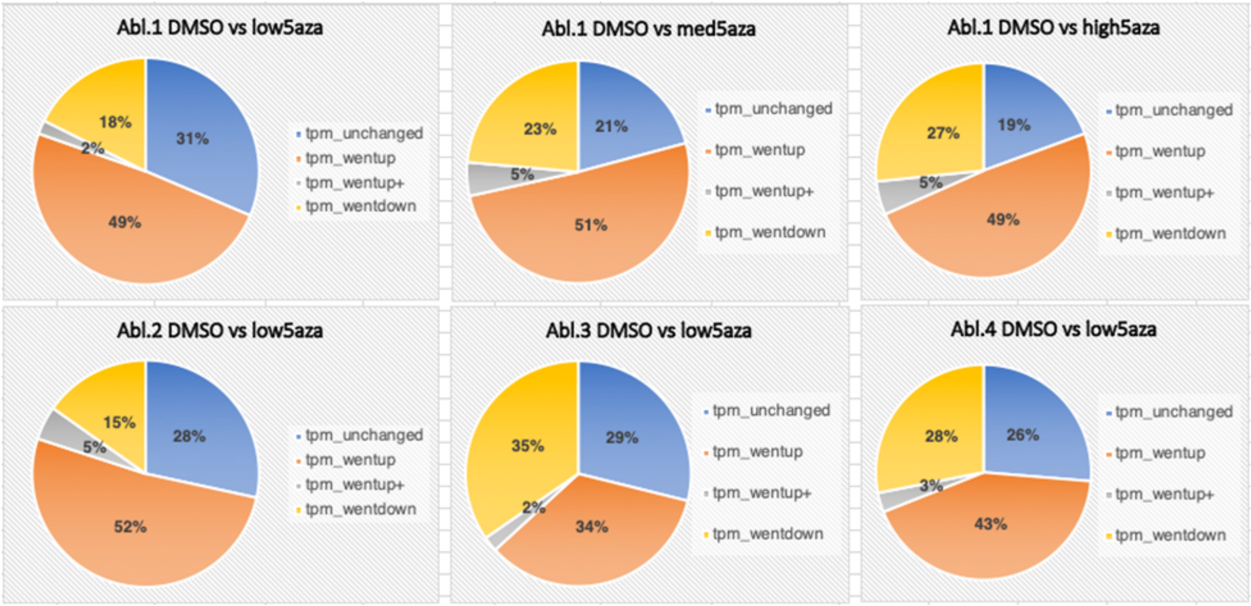

**Supplemental Figure S12. Change in RNA abundance for genes showing differential AI. Related to Figure 4C and 4F.**
Percentage of genes showing changes in RNA abundance for genes showing significant differential AI between control and treated condition are shown as a pie charts. Complete data in Suppl. Table S8.

tpm unchanged: genes that do not change TPM ($P_{adj}$ >0.05)
tpm wentup: genes that increase TPM by more than 2-fold after drug treatment ($P_{adj}$<0.05)
tpm wentup+: genes that increase TPM by more than 4-fold after drug treatment ($P_{adj}$ <0.05)
tpm wentup: genes that decreases TPM below 2-fold after drug treatment ($P_{adj}$<0.05)

(A)

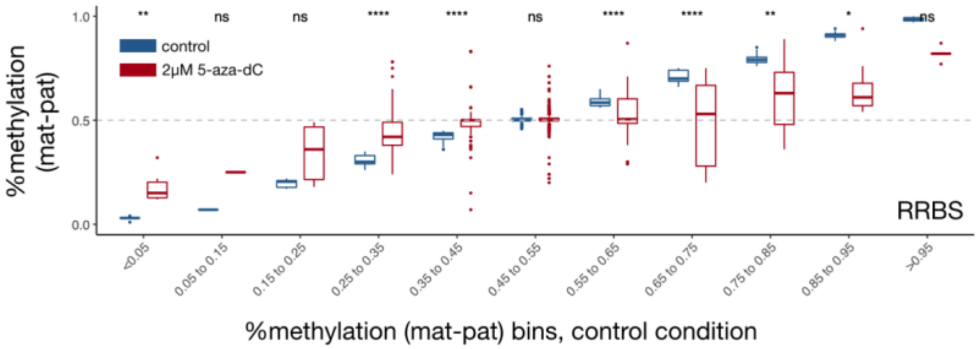

(B)

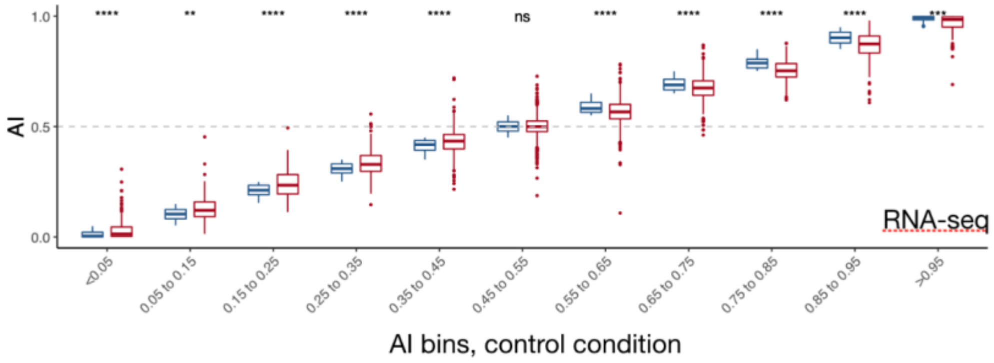

**Supplemental Figure S13. Allele-specific impact on transcriptome-wide gene regulatory landscape in Abl.1 cells. Abl.1 cells were exposed for 2 days to 2µM 5-aza-dC in 1% DMSO or to only 1% DMSO control. Related to Figure 4D-E.**

(A) [% methylation (maternal – paternal)] or diff denotes difference in percentage methylation of maternal and paternal alleles measured using reduced representation bisulfite sequencing (RRBS). Percent methylation for maternal allele was calculated by taking the ratio of methylated reads for maternal allele of a gene to total reads for that gene. Diff value range from -1 to +1 and was normalised from 0 to 1, shown on Y-axis. Assessed regions were first binned by their diff values in control condition. Boxplots were then generated for each condition separately, *blue*: in control condition (1% DMSO) and *red*: in 2 µM 5-aza-dC. Mean comparisons were made, and P-values were calculated for all control and treated groups using Wilcoxon test. The extreme bins showed non-significant (ns) difference due to insufficient data in those bins.

(B) allelic imbalance was measured using RNA-seq. Data is same as shown in Fig. 4E for RNA-seq. Mean comparisons were made, and P-values were calculated for all control (*blue* boxplots) and treated (*red* boxplots) groups using Wilcoxon test.

| clone | Abl.2 | | | Abl.3 | | | Abl.4 | | |
|---|---|---|---|---|---|---|---|---|---|
| 5-aza-dC conc | 0 | 0.1 | 0.2 | 0 | 0.1 | 0.2 | 0 | 0.1 | 0.2 |
| day 0 | 94% | | | 96% | | | 95% | | |
| day 1 | 93% | 72% | 77% | 93% | 67% | 55% | 95% | 77% | 81% |
| day 2 | 87% | 41% | 24% | 84% | 30% | 11% | 84% | 27% | 31% |

**Supplemental Figure S14. Cell viability of clones, Abl.2, Abl.3 and Abl.4 in 5-aza-dC. Related to Figure 4F.** Percent of live cells on each day for concentrations tested (in µM) are shown. 0µM refers to cells grown in presence of only drug vehicle of 1% DMSO.

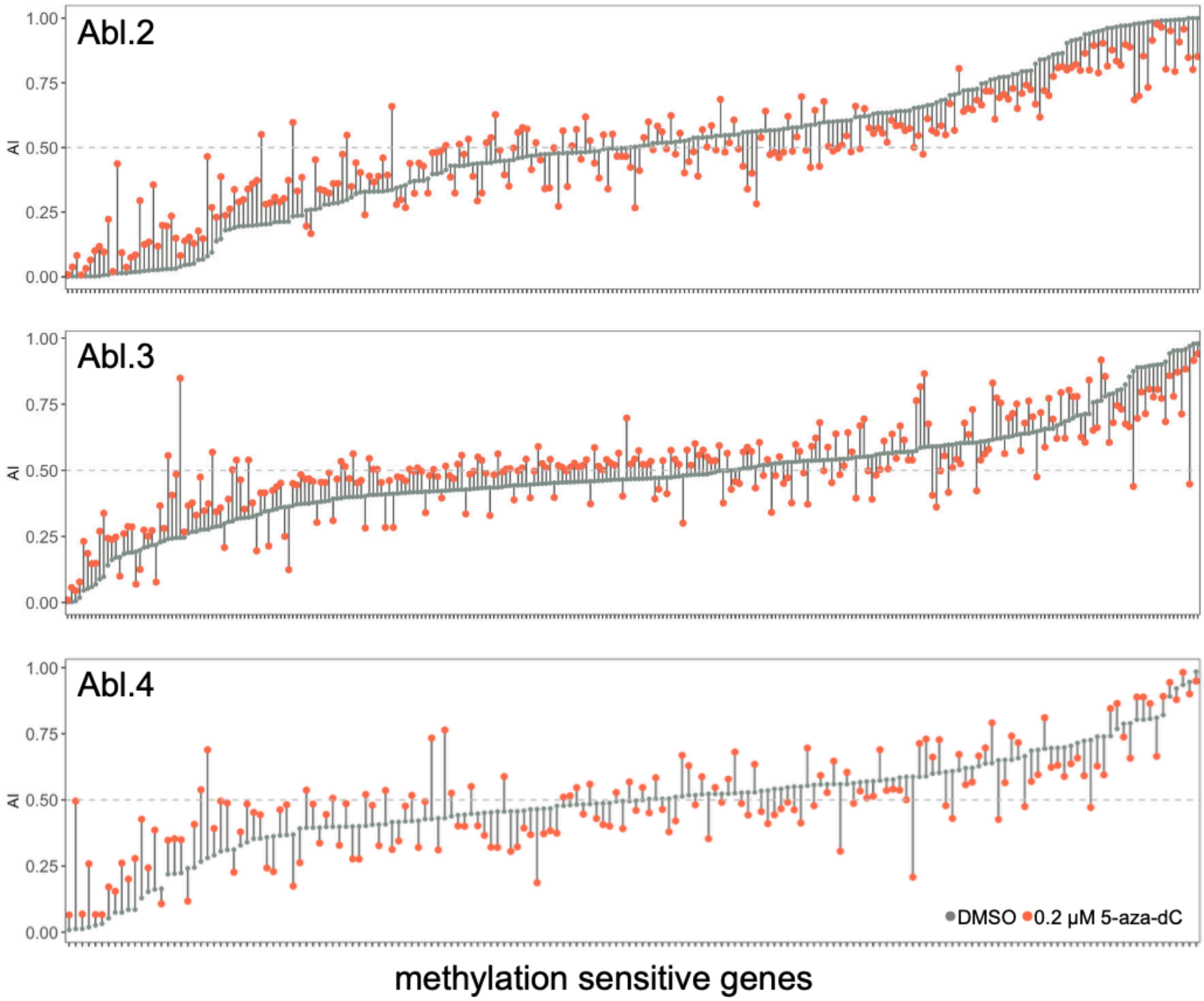

**Supplemental Figure S15. Autosomal genes showing significant differential AI between DMSO and 0.2µM 5-aza-dC in clones Abl.2, Abl.3 and Abl.4. Related to Figure 4F.**
*Grey circles* show AI in control and *red circles* shows AI after treatment. Genes are ordered by their AI in 1% DMSO. See **Suppl. Table S7** for complete data.

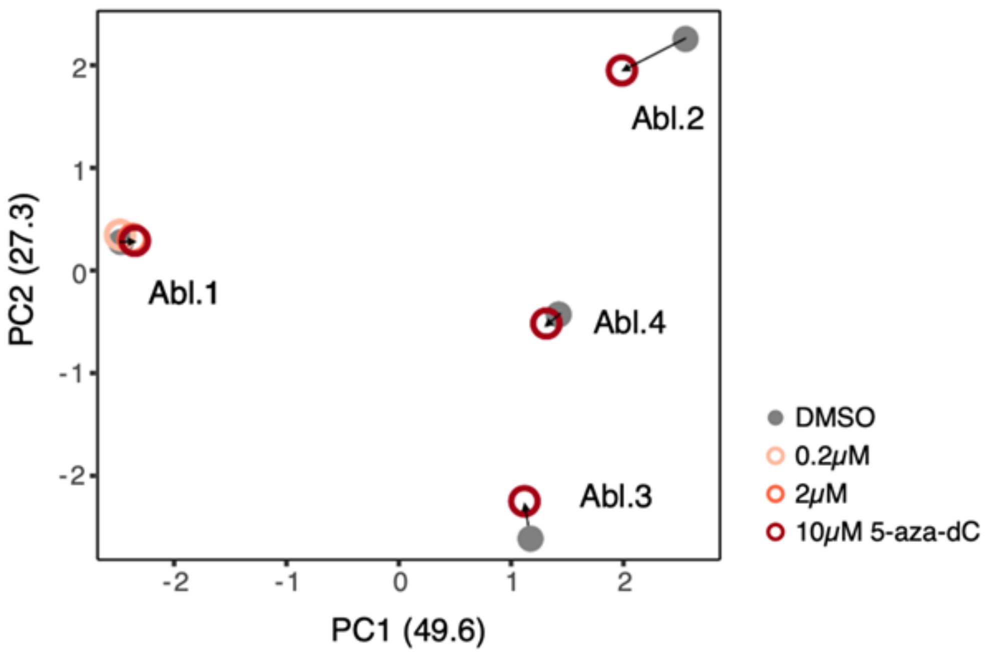

**Supplemental Figure S16. PCA of all methylation-sensitive genes in all Abelson clones before and after 5-aza-dC treatment. Related to Figure 4F.**
Principal component analysis of AI of 677 methylation sensitive genes in all clonal cell lines, Abl.1, Abl.2, Abl.3 and Abl.4 in 1% DMSO (*grey*) moving closer to each other in presence of 5-aza-dC (*colored circles*).

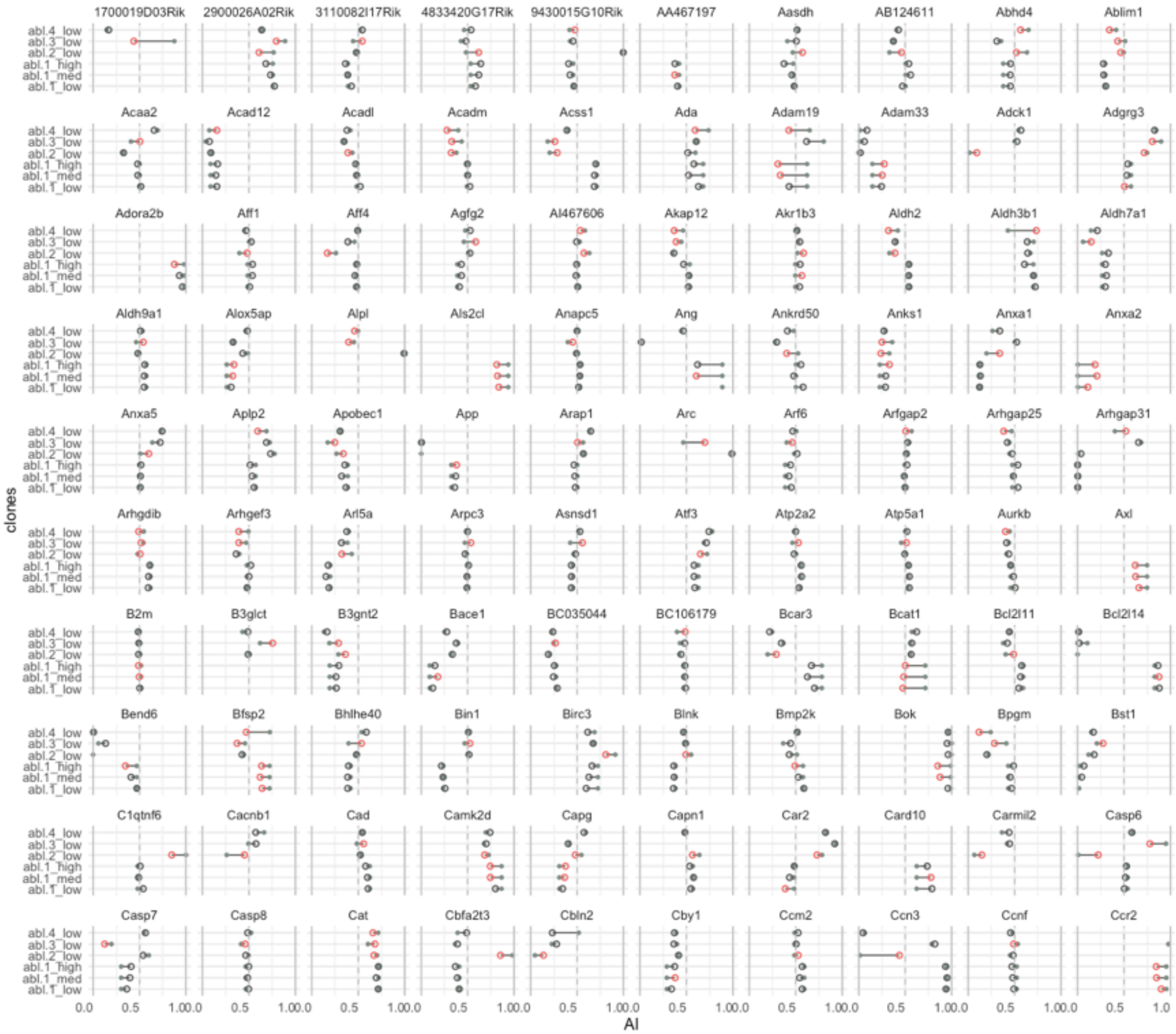

Supplemental Figure S17 (cont’d on next page)

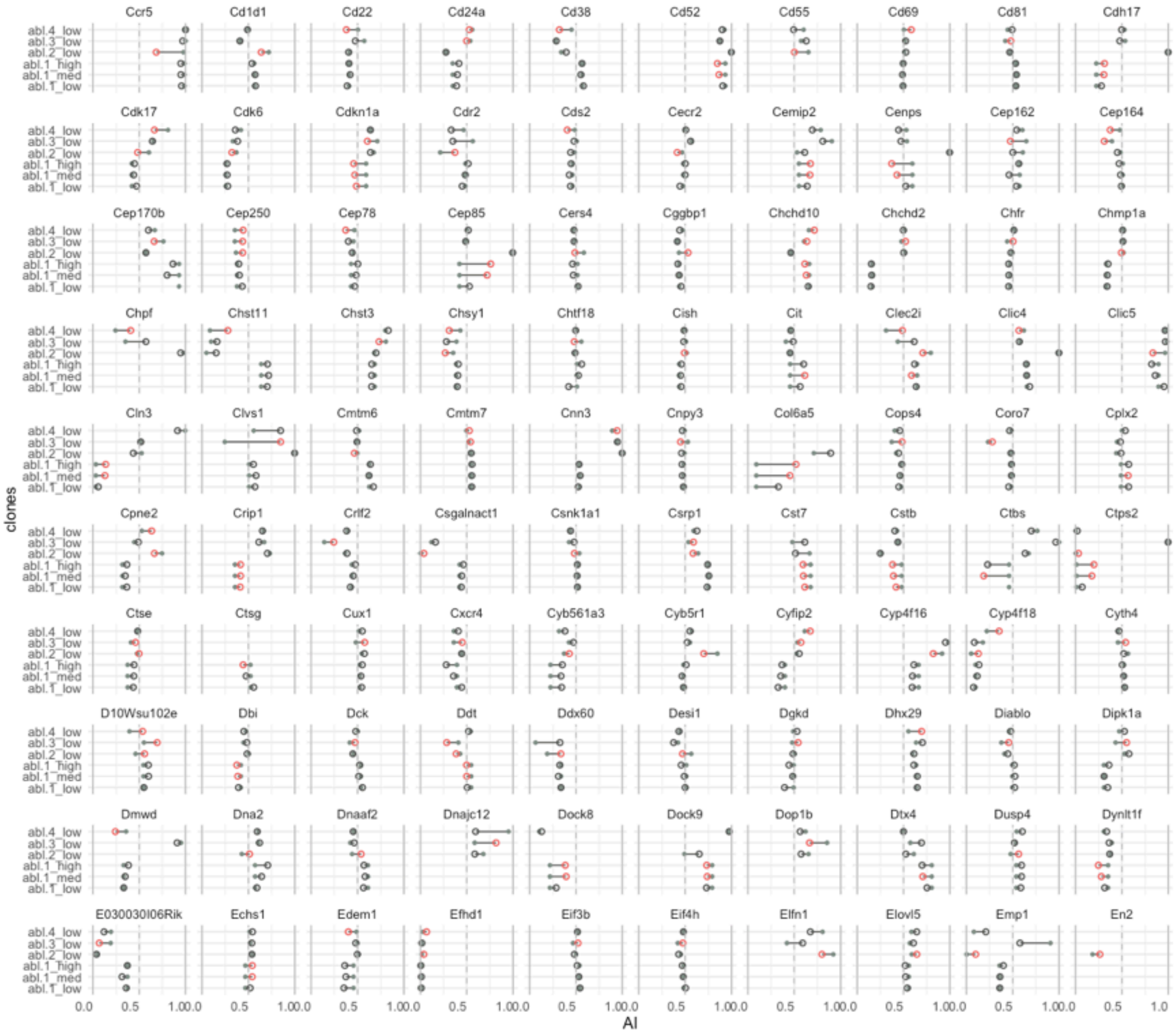

Supplemental Figure S17 (cont'd)

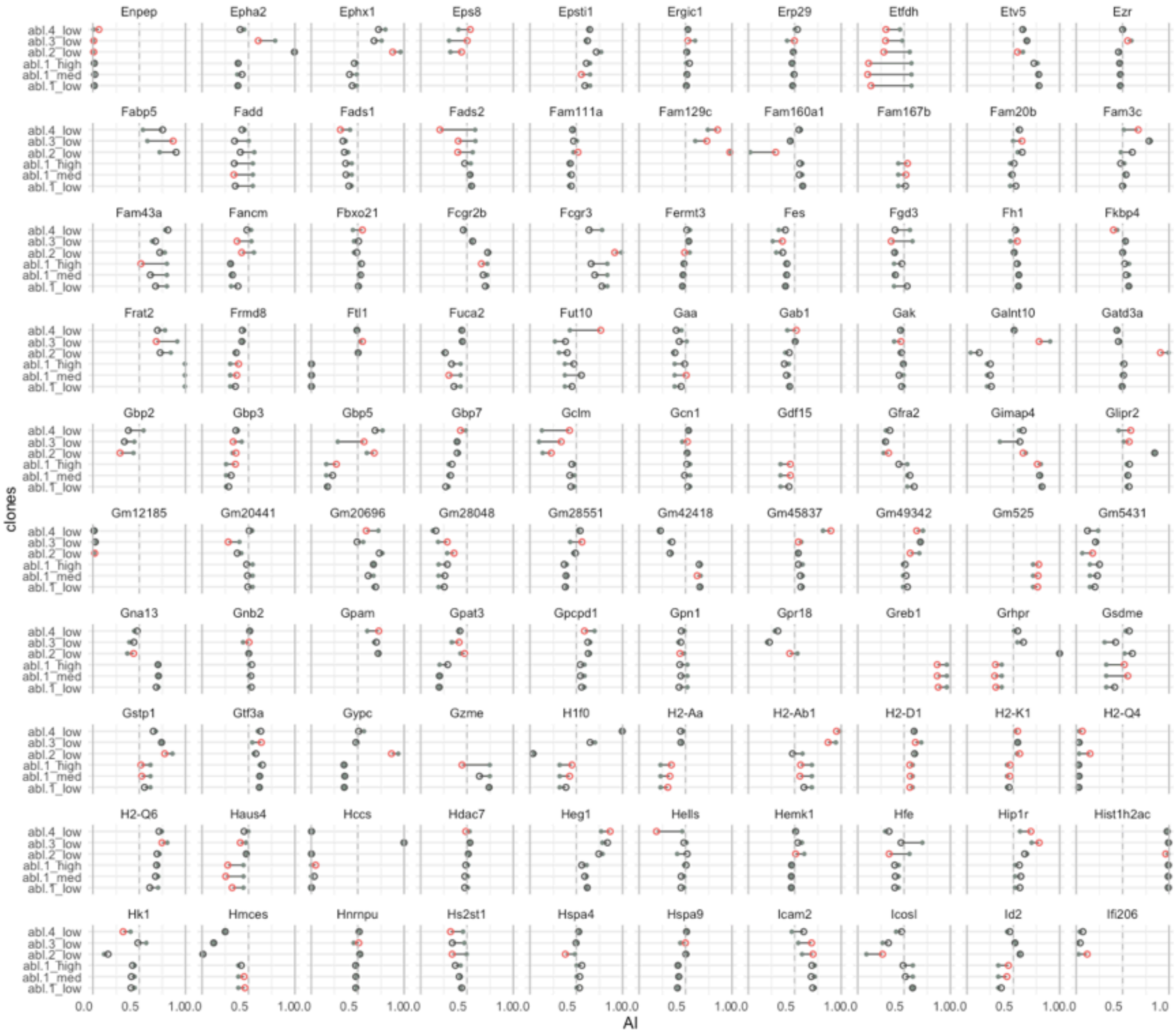

Supplemental Figure S17 (cont'd)

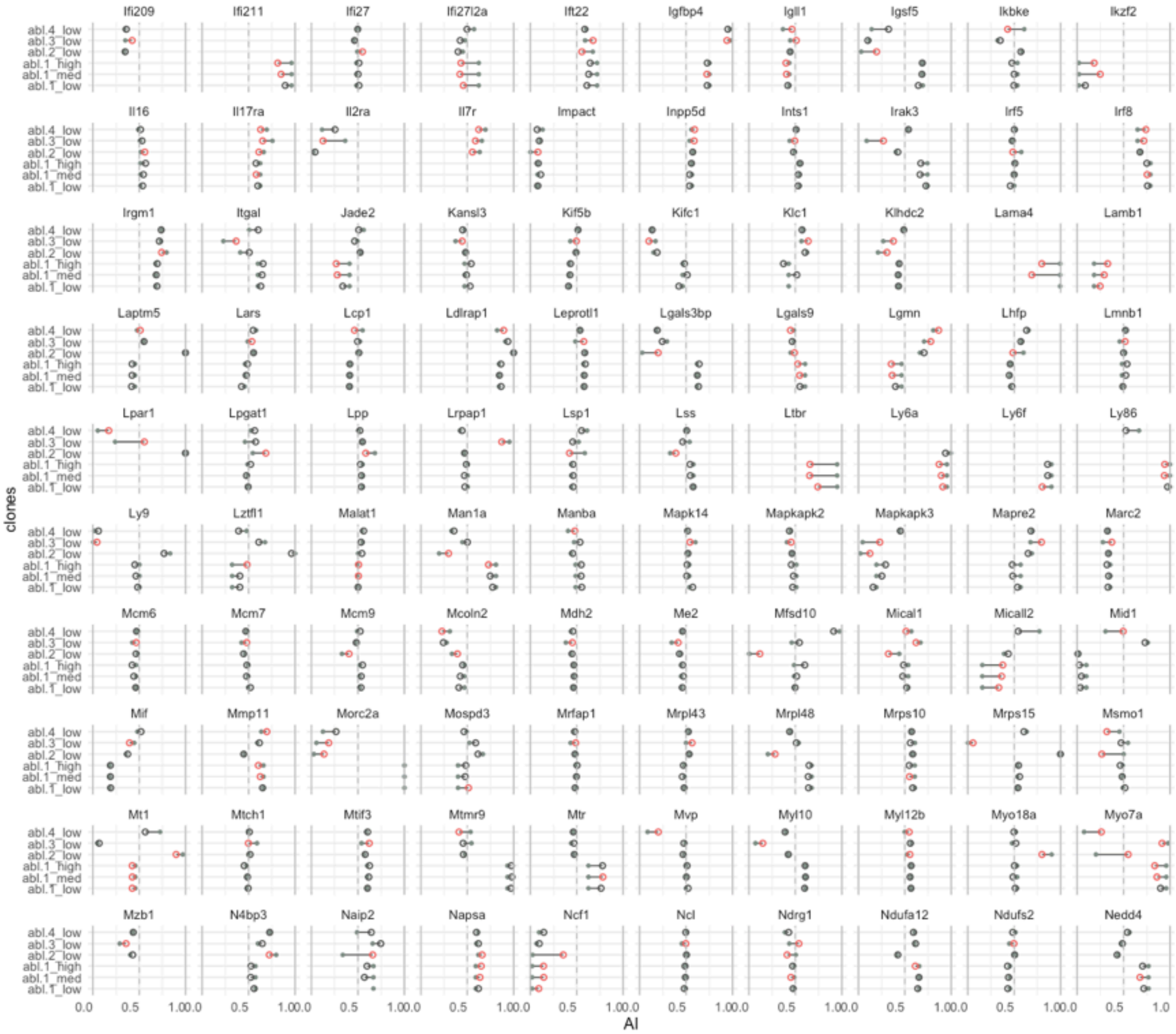

Supplemental Figure S17 (cont’d)

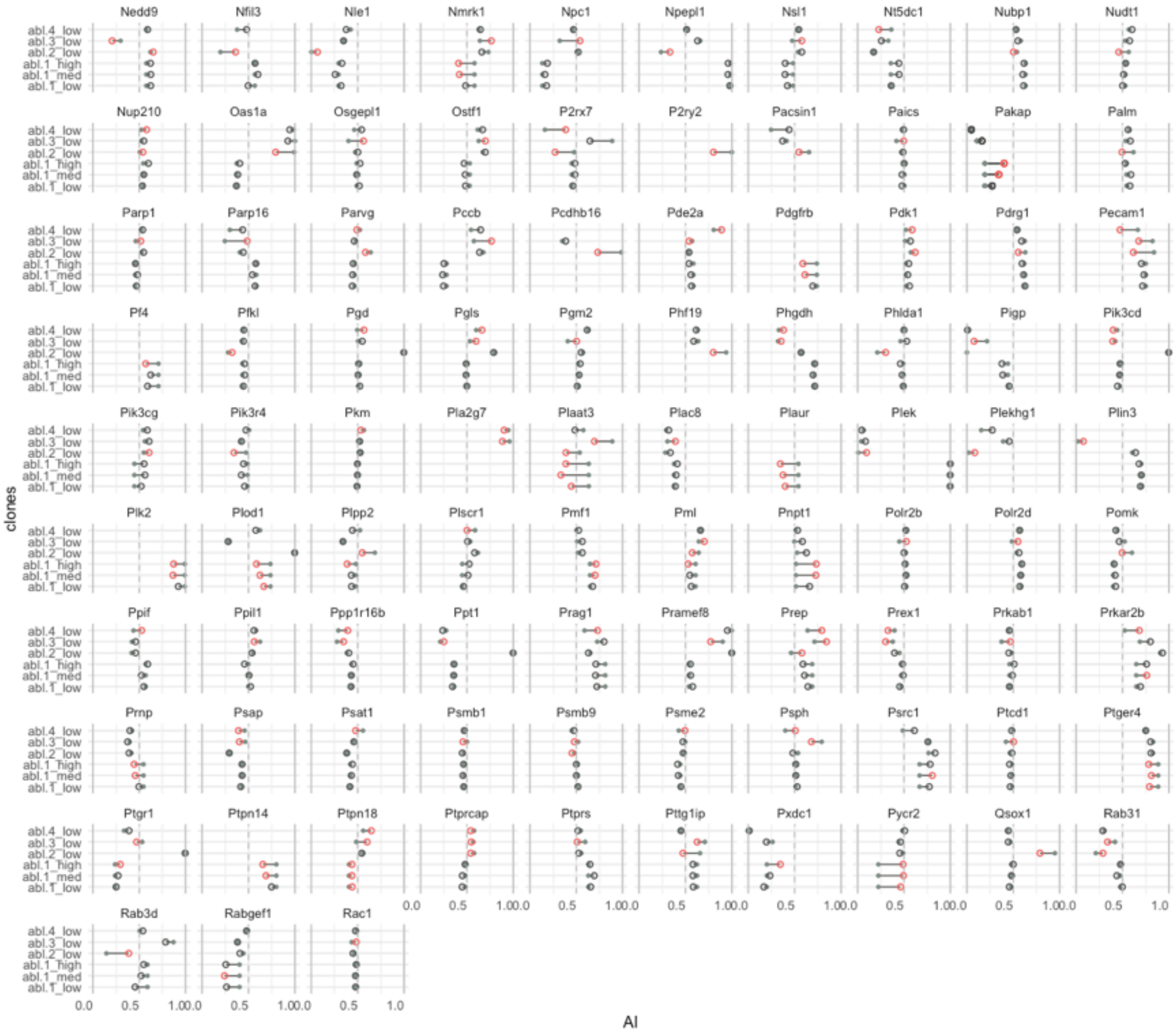

Supplemental Figure S17 (cont’d)

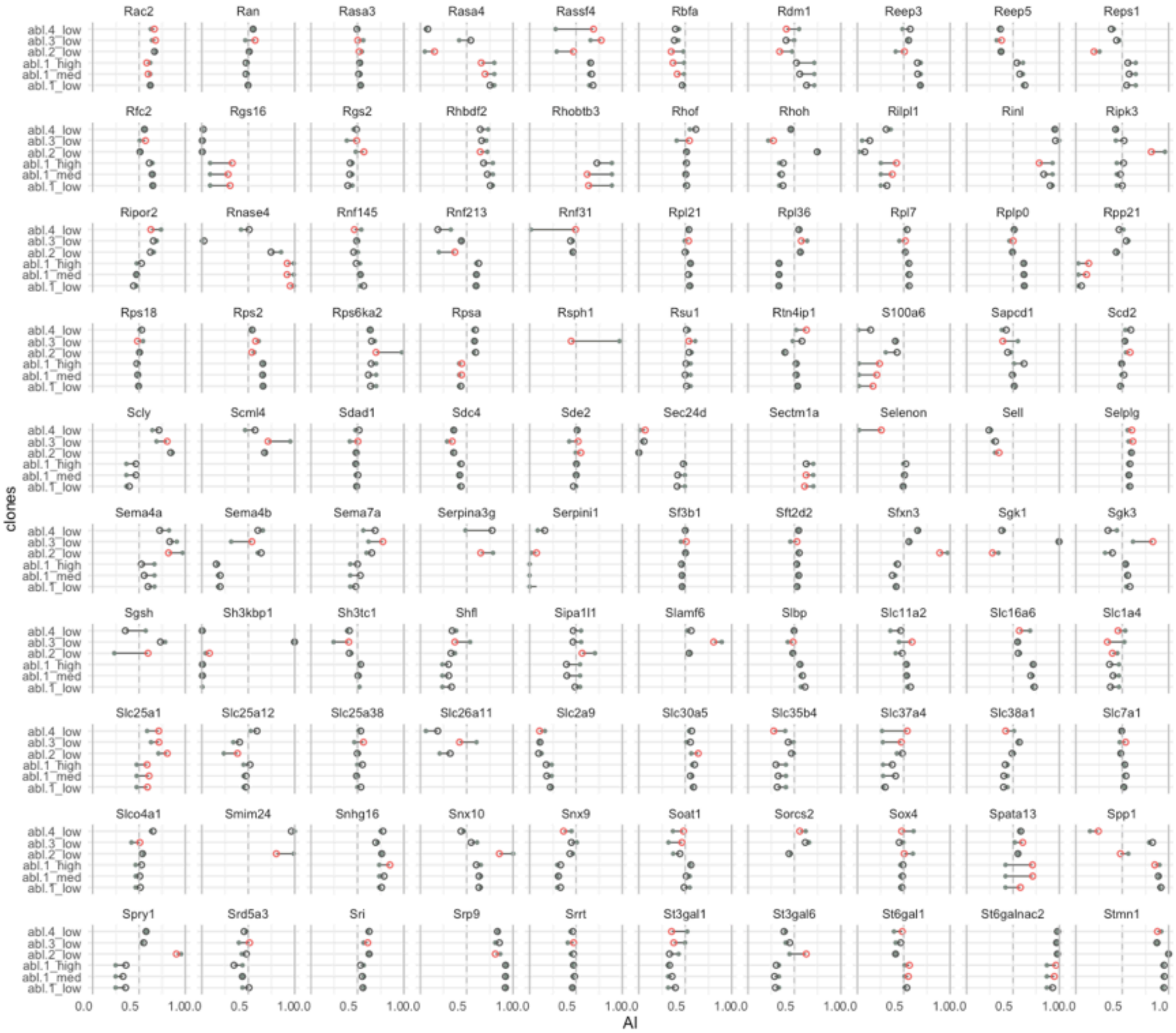

Supplemental Figure S17 (cont'd)

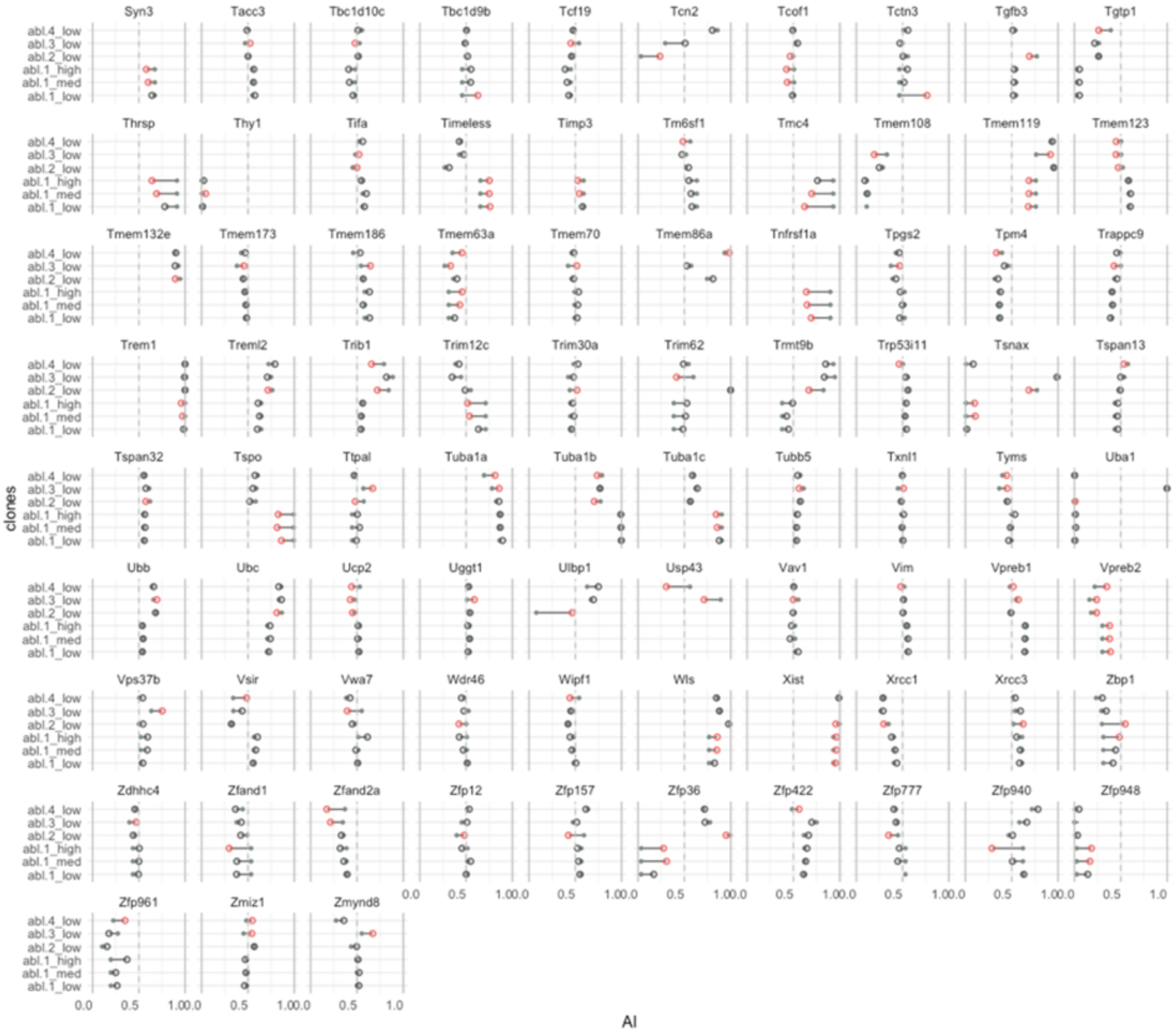

**Supplemental Figure S17. 677 genes showing significant differential AI in any of the four assessed Abelson clones. Related to Figure 4F.**

AI before and after 5-aza-dC treatment in any of the clones, Abl.1 (0.2µM, 2µM, 10µM), Abl.2 (0.2µM), Abl.3 (0.2µM), and Abl.4 (0.2µM) is shown. AI in control is shown as *grey solid* circles and in 5-aza-dC is shown as *open* circles. *Open* circles are colored *red* if the shift in AI is significant in that clone, else are colored *black*.

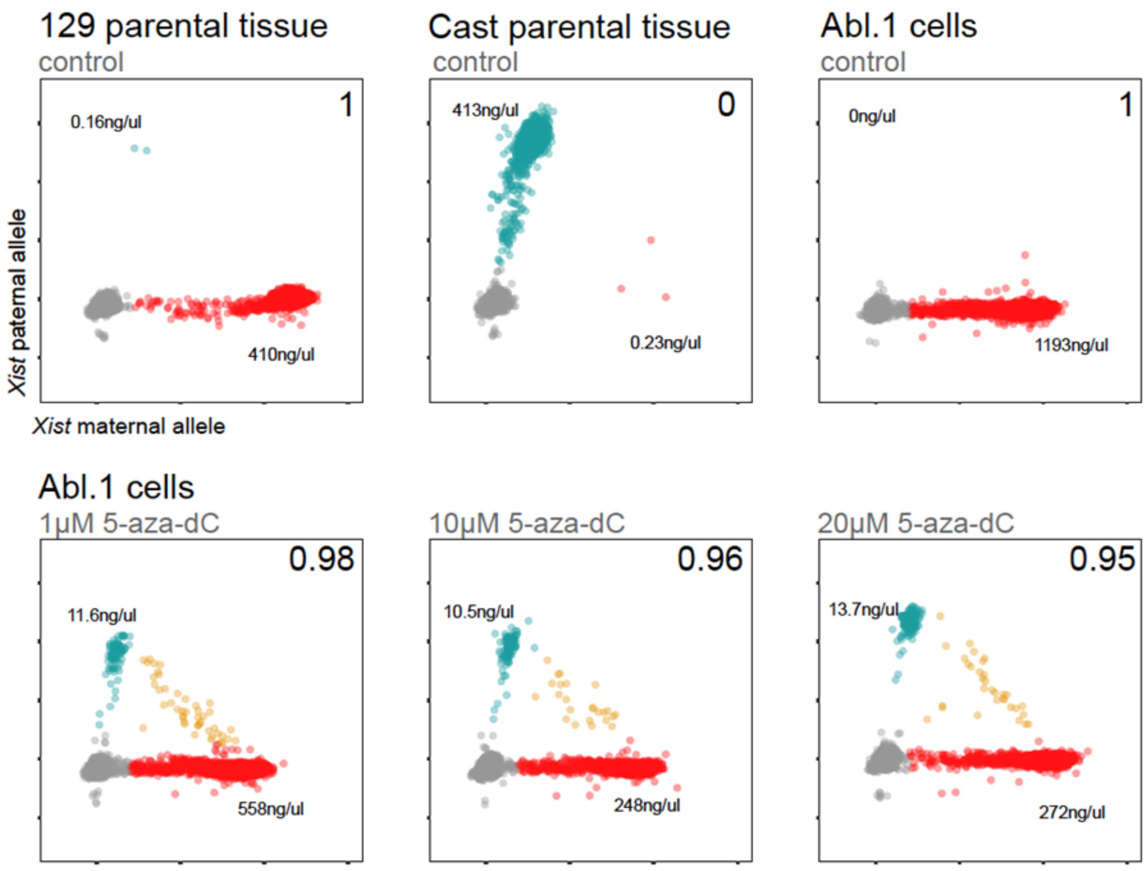

**Supplemental Figure S18. *Xist* allele imbalance in Abl.1 cells on 5-aza-dC treatment with ddPCR. Related to Figure 2A and B.**
Scatterplots for 20,000 droplets targeting *Xist* in control untreated samples (*upper panel*) and 5-aza-dC treated Abl.1 cells (*lower panel*). cDNA samples from day 7 of screening were assessed using ddPCR with allele-specific fluorescent probes. *Grey*: empty droplets; *blue*: droplets with Cast allele amplified; *red*: droplets with 129 allele amplified. Ratio of red:blue droplets (AI) is written on the upper right corner of each plot. Next to red and blue droplets are the copies/µl measured for 129 and Cast *Xist* allele.

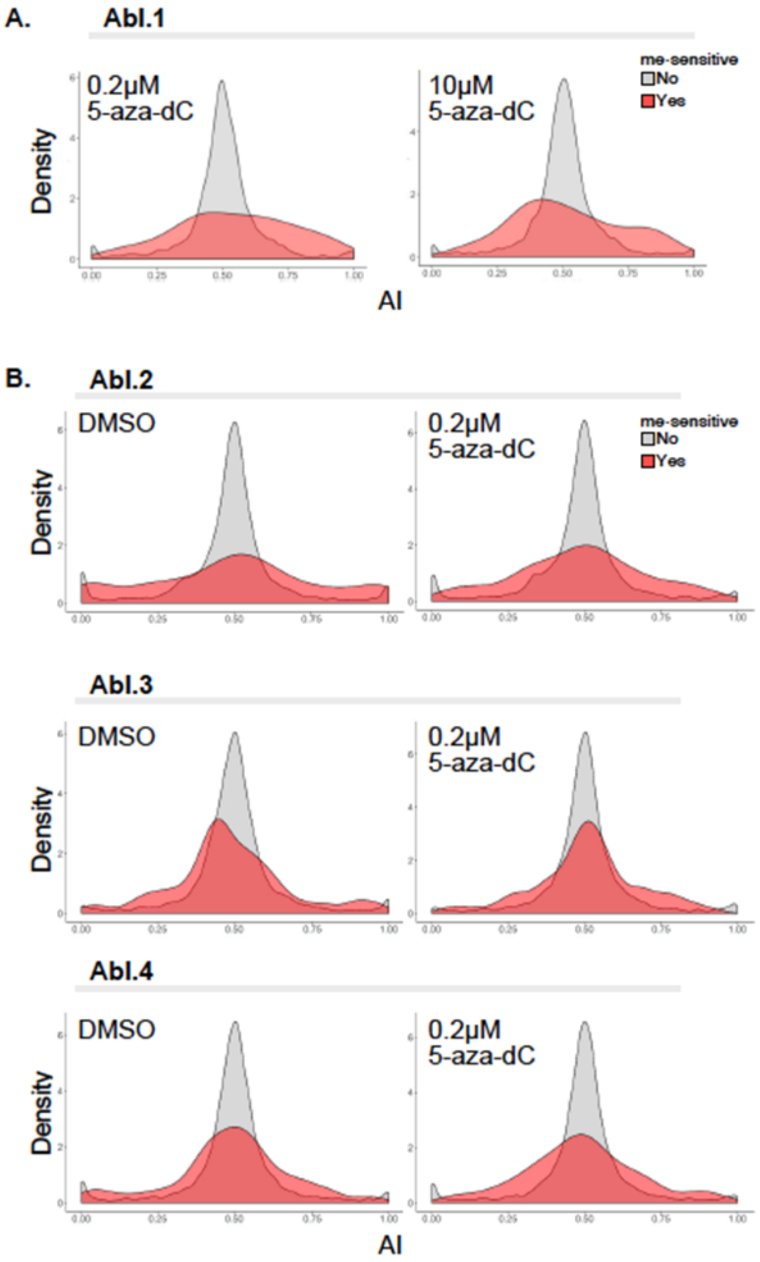

**Supplemental Figure S19. Density plots for distribution of AI values in control and 5-aza-dC treatment for Abelson clones. Related to Figure 4B.**

Density plots for distribution of AI values of genes with no significant changes in AI (*grey*) and with changes (*red*) on 5-aza-dC treatment. Note that grey and red areas are plotted to be equal. (A.) in Abl.1 cells. Left: distribution in 0.2µM 5-aza-dC; right: distribution in 10 µM 5-aza-dC (B.) in Abl.2, Abl.3 and Abl.4 cells. Left: distribution in 1% DMSO; right: distribution in 0.2 µM 5-aza-dC

**Supplemental Table S1. Description of readout genes used in Screen-seq, AI in clone Abl.1 and Abl.2. Related to Fig. 1.**
*red* - MAE genes with maternal bias, *blue* – MAE genes with paternal bias, *yellow* – Biallelic genes, *no color* – control genes, imprinted or on X-chromosome, in clone Abl.1. Data from Nag *et a*l 2015 (G3). Primers for these genes are in **Suppl. Table S2**.

| Readout genes | chr | Class (based on Abl.1) | Multiplex assay | Allelic bias (maternal AI:paternal AI) | | RNA abundance (TPM) | |
|---|---|---|---|---|---|---|---|
| | | | | Abl.1 | Abl.2 | Abl.1 | Abl.2 |
| *Dlc1* | chr08 | MAE_Cast | UMI | 0.1:0.9 | 1.0:0.0 | 7.89 | 0.33 |
| *Fam217b* | chr02 | MAE_Cast | Non-UMI | 0.0:1.0 | 1.0:0.0 | 11.36 | 0.28 |
| *Spire1* | chr18 | MAE_Cast | Non-UMI | 0.1:0.9 | 1.0:0.0 | 6.16 | 0.21 |
| *Col6a5* | chr09 | MAE_Cast | Non-UMI | 0.0:1.0 | 1.0:0.0 | 15.39 | 0.31 |
| *Pea15a* | chr01 | MAE_Cast | Non-UMI | 0.0:1.0 | 0.8:0.2 | 41.46 | 4.5 |
| *H2-Ea-ps* | chr17 | MAE_Cast | Non-UMI | 0.0:1.0 | 1.0:0.0 | 74.43 | 0.42 |
| *Sash1* | chr10 | MAE_Cast | Non-UMI | 0.0:1.0 | 1.0:0.0 | 15.27 | 0.29 |
| *Dnajc12* | chr10 | MAE_Cast | Both | 0.0:1.0 | 0.8:0.2 | 5.53 | 2.75 |
| *Adamtsl4* | chr03 | MAE_Cast | Both | 0.0:1.0 | 1.0:0.0 | 13.1 | 0.99 |
| *3830406C13Rik* | chr14 | MAE_129 | UMI | 1.0:0.0 | 0.0:1.0 | 13.02 | 3.6 |
| *Tpst1* | chr05 | MAE_129 | UMI | 0.9:0.1 | 0.4:0.6 | 8.34 | 24.79 |
| *Afap1* | chr05 | MAE_129 | Non-UMI | 1.0:0.0 | 0.0:1.0 | 2.43 | 3.81 |
| *Ncam2* | chr16 | MAE_129 | Non-UMI | 1.0:0.0 | 0.0:1.0 | 2.95 | 2.9 |
| *Smtnl2* | chr11 | MAE_129 | Both | 1.0:0.0 | 33:67 | 9.02 | 12.5 |
| *Adnp2* | chr18 | MAE_129 | Both | 1.0:0.0 | 0.0:1.0 | 17.8 | 8.45 |
| *Slc25a37* | chr14 | BAE | UMI | 0.4:0.6 | 0.5:0.5 | 73.54 | 30.91 |
| *Ccnj* | chr19 | BAE | Non-UMI | 0.5:0.5 | 0.3:0.7 | 8.35 | 4.32 |
| *Hdlbp* | chr01 | BAE | Non-UMI | 0.6:0.4 | 0.6:0.4 | 192.14 | 40.07 |
| *Casd1* | chr06 | 129_Imprinted | Non-UMI | 1.0:0.0 | 0.3:0.7 | 18.28 | 12.13 |
| *Pola1* | chrX | X-linked | Non-UMI | 0.0:1.0 | 0.0:1.0 | 56.71 | 44.26 |
| *Phka2* | chrX | X-linked | Non-UMI | 1.0:0.0 | 0.0:1.0 | 44.72 | 12.71 |
| *Mecp2* | chrX | X-linked | Non-UMI | 1.0:0.0 | 0.0:1.0 | 22.42 | 12.71 |
| *Gprasp1* | chrX | X-linked | UMI | 0.0:1.0 | 0.0:1.0 | 10.12 | 3.63 |

**Supplemental Table S6. Datasets generated and analyzed in this study. Related to Fig. 4.**

| Data | Sample | Rep | # Fragments seq | Seq type | GEO ID |
|---|---|---|---|---|---|
| RNAseq | 1. Abl.1 – DMSO | 1 | 43 514 706 | SE75 | GSE144005 |
| | 2. Abl.1 – DMSO | 2 | 49 967 582 | SE75 | |
| | 3. Abl.1 – 0.2µM 5-aza-dC | 1 | 43 249 221 | SE75 | |
| | 4. Abl.1 – 0.2µM 5-aza-dC | 2 | 21 371 037 | SE75 | |
| | 5. Abl.1 – 2µM 5-aza-dC | 1 | 21 371 037 | SE75 | |
| | 6. Abl.1 – 2µM 5-aza-dC | 2 | 30 374 302 | SE75 | |
| | 7. Abl.1 – 2µM 5-aza-dC | 3 | 45 291 427 | SE75 | |
| | 8. Abl.1 – 2µM 5-aza-dC | 4 | 40 853 428 | SE75 | |
| | 9. Abl.1 – 2µM 5-aza-dC | 5 | 44 156 766 | SE75 | |
| | 10. Abl.1 – 10µM 5-aza-dC | 1 | 42 751 147 | SE75 | |
| | 11. Abl.1 – 10µM 5-aza-dC | 2 | 38 845 075 | SE75 | |
| | 12. Abl.2 – DMSO | 1 | 34 072 961 | PE150 | |
| | 13. Abl.2 – DMSO | 2 | 42 493 030 | PE150 | |
| | 14. Abl.2 – 0.2µM 5-aza-dC | 1 | 39 133 069 | PE150 | |
| | 15. Abl.2 – 0.2µM 5-aza-dC | 2 | 59 234 002 | PE150 | |
| | 16. Abl.3 – DMSO | 1 | 30 128 487 | PE150 | |
| | 17. Abl.3 – DMSO | 2 | 29 282 670 | PE150 | |
| | 18. Abl.3 – 0.2µM 5-aza-dC | 1 | 36 931 828 | PE150 | |
| | 19. Abl.3 – 0.2µM 5-aza-dC | 2 | 35 273 378 | PE150 | |
| | 20. Abl.4 – DMSO | 1 | 30 575 079 | PE150 | |
| | 21. Abl.4 – DMSO | 2 | 34 217 388 | PE150 | |
| | 22. Abl.4 – 0.2µM 5-aza-dC | 1 | 36 225 623 | PE150 | |
| | 23. Abl.4 – 0.2µM 5-aza-dC | 2 | 39 678 430 | PE150 | |
| RRBS | 1. Abl.1 – DMSO | 1 | 8 874 755 | PE100 | GSE144006 |
| | 2. Abl.1 – DMSO | 2 | 5 311 764 | PE100 | |
| | 3. Abl.1 – 0.2µM 5-aza-dC | 1 | 9 543 598 | PE100 | |
| | 4. Abl.1 – 0.2µM 5-aza-dC | 2 | 8 746 138 | PE100 | |
| | 5. Abl.1 – 2µM 5-aza-dC | 1 | 9 073 154 | PE100 | |
| | 6. Abl.1 – 2µM 5-aza-dC | 2 | 9 671 719 | PE100 | |
| | 7. Abl.1 – 2µM 5-aza-dC | 3 | 10 339 090 | PE100 | |
| | 8. Abl.1 - 10µM 5-aza-dC | 1 | 8 701 011 | PE100 | |
| | 9. Abl.1 - 10µM 5-aza-dC | 2 | 5 797 257 | PE100 | |

The following tables are in Main Supplemental Tables:

**Suppl. Table S2**
Readout genes: Primers for Screen-seq, ddPCR and RT-qPCR. Related to Figure 1.

**Suppl. Table S3**
Drug library tested on Abl.1 clonal cell line. Related to Figure 1.

**Suppl. Table S4**
Readout genes: All drug screen hits. Related to Figure 1.

**Suppl. Table S5**
Complete Screen-seq results. Related to Figure 1.

**Suppl. Table S7**
677 methylation sensitive genes in all Abelson clones. Related to Figure 4.

**Suppl. Table S11**
1767 genes with significant differential AI between Abelson clones.

**Suppl. Table S12**
346 methylation sensitive genes with significant differential AI between Abelson clones.

The following supplemental tables are in separate files:

**Suppl. Table S8**
Change in RNA abundance for methylation sensitive genes in all Abelson clones. Related to Figure 4.

**Suppl. Table S9**
Complete genome-wide AI measurement in all Abelson clones with RNA-seq on 5-aza-dC exposure. Related to Figure 4.

**Suppl. Table S10**
Complete genome-wide AI measurement in Abl.1 clone with RRBS on 5-aza-dC exposure. Related to Figure 4.
